# Kinetics and dynamics of single-molecule multivalent interactions revealed by plasmon-enhanced fluorescence

**DOI:** 10.1101/2023.12.08.570798

**Authors:** Kasper R. Okholm, Sjoerd W. Nooteboom, Vincenzo Lamberti, Swayandipta Dey, Peter Zijlstra, Duncan S. Sutherland

**Affiliations:** Interdisciplinary Nanoscience Center, Aarhus University, Gustav Wieds Vej 14, Aarhus C, 8000, Denmark; The Centre for Cellular Signal Patterns (CELLPAT), Gustav Wieds Vej 14, Aarhus C, 8000, Denmark; Eindhoven University of Technology, Department of Applied Physics and Science Education and Institute for Complex Molecular Systems, 5600 MB Eindhoven, The Netherlands

## Abstract

Multivalency as an interaction principle is widely utilized in nature. It enables specific and strong binding by multiple weak interactions through enhanced avidity and is a core process in immune recognition and cellular signaling and a current concept in drug design. Rapid binding and unbinding of monovalent constituent interactions during multivalent binding creates dynamics that require a single-molecule approach to be studied. Here, we use the high signals from plasmon enhanced fluorescence of nanoparticles to extract binding kinetics and dynamics of multivalent interactions on the single-molecule level and in real-time. We study mono-, bi-and trivalent binding interactions using a DNA Holliday Junction as a model construct with programmable valency. Furthermore, we introduce a model framework for binding kinetics that involves the binding restriction during multivalent interactions to take into account the structural conformation of multivalent molecules allowing quantitative comparison. We used this approach to explore how length and flexibility of the DNA ligands affect binding restriction and binding strength, where overall binding strength decreased with spacer length. For trivalent systems increasing spacer length was found to activate binding in the trivalent state giving insight into the design of multivalent drug or targeting moieties. Interestingly we could exploit the rapidly decaying near fields of the plasmon that induce a strong dependence of the signal to position of the fluorophore to observe binding dynamics during single multivalent binding events.

## Introduction

In nature, the principle of multivalency enables specific and strong, but still reversible, binding between receptors and ligands^1–3^. Multivalency involves binding of multiple reversible chemical interactions between two units where each interaction is relatively weak but give rise to dramatically enhanced binding strengths due to enhanced avidity also known as functional affinity^4^. The interaction of viruses with hosts cells is an important example that is mediated through multiple weak bonds, leading to uptake by the host cells via strong multivalent binding^1^. These multivalent interactions can occur between a single receptor-ligand pair where multiple binding motifs are involved^5^, between a multivalent object and a surface (e.g. virus-host cell interaction^6^ or immune macromolecular recognition of a pathogen surface^7^) or between two cell surfaces leading to adhesion or signaling^8^. Multivalency has been subject to strong research interest for more than 25 years since the seminal paper by Whitesides et al^1^.

The perspectives of studying multivalency are manyfold. One is to fundamentally understand multivalent interactions in order to give insight into different natural molecular and cellular processes. Another perspective is to use the underlying principles of multivalent binding in drug design^9–11^, molecular imaging agents^12^, in supramolecular chemistry^2,13^, and for functional materials^14,15^. Various models on the nature of multivalent interactions have been proposed and used to explain different multivalent systems in terms of their thermodynamic and kinetic properties^1,16–19^. Though these models have had great success at explaining fundamental concepts, there is still far to go to understand multivalent binding in terms of molecular structure, flexibility and steric effects in order to unleash the potential of multivalency in drug design and to understand the mechanism of drug action during blocking of multivalent drug targets^20^.

Multivalent binding is often measured and evaluated based on ensemble-averaged techniques and assays. Common examples are surface plasmon resonance, ELISA, ICT50, and bilayer interferometry^2^. These techniques allow easy quantification of binding constants and avidity enhancement. However, multivalent interactions are inherently heterogenous and require a single-molecule approach to understand the underlying dynamic behavior during binding that drives the avidity and can lead to surface mobility and walking^21^. A range of single-molecule approaches have been used on multivalent systems e.g. single-molecule fluorescence^22^, force spectroscopy using AFM^23,24^ and nanopores^25^. These techniques have been used to study the binding kinetics of multivalent systems but none of these techniques have been able to capture the dynamics of a multivalent interaction during a binding event.

Here, we present a method that enables extraction of binding kinetics while also providing direct observations of binding dynamics in multivalent interactions on the single-molecule level. We demonstrate the approach to study mono-, bi-, and trivalent binding and the principles for macromolecular design. Our approach, termed plasmon enhanced fluorescence of multivalent systems at the single-molecule level (PEFOMS), makes use of plasmon enhanced fluorescence from gold nanoparticles^26^ that provides superior signals of binding events^27^. Importantly, the rapidly decaying optical fields near the particles allow us to directly observe binding dynamics during individual multivalent ligand-receptor binding events. We use a DNA based interaction framework of DNA coated nanoparticles and 4-way junctions as a multivalent ligand-receptor system with programmable valency and binding strength. Here we apply the single-molecule approach to study avidity and observe complex dynamics during single binding events for multivalent systems and compare to an extended step-binding model taking into account macromolecular conformation. Our approach can provide insight into biologically relevant multivalent interactions and inform the design of synthetic multivalent macromolecules.

## Results and discussion

### Plasmon enhanced fluorescence from single molecules binding to gold nanorods

We use PEFOMS to measure the bound-state lifetimes of fluorescently labeled molecules. The principle is illustrated in figure 1 and adopts the framework used in DNA based points accumulation for imaging in nanoscale topography (DNA-PAINT)^28,29^. In our approach we use DNA coated gold nanorods immobilized on a coverslip and imaged in a total internal reflection (TIRF) based microscope ^30,31^. Under illumination gold nanorods support localized surface plasmon resonances (LSPR) from the collective oscillations of electrons in the particle^32^. At the plasmon resonance the electric field is strongly enhanced in the vicinity of the nanorod, leading to distance dependent excitation enhancement. In addition, the coupling between the dye and the particle results in modification of the radiative and non-radiative decay rates, leading to a distance-dependent emission enhancement ^27,33–35^. Using plasmon enhanced fluorescence on DNA coated gold nanorods it is possible to track binding events of dye labelled DNA probes and quantify binding kinetics^30^.The nanorods are functionalized with 30nt long thiolated DNA strands here termed *receptor strands*. The end-segment carries a specific sequence that a complementary DNA sequence, termed *ligand DNA*, can transiently bind to depending on the sequence and length of complementarity thereby representing a ligand-receptor interaction. The receptor DNA was attached using a freeze-thawing approach to maximize the density of DNA on the gold particle surface^36^. The average receptor density was 1.6 moles/*cm^2^* corresponding to a distance of 3.5 nm between receptor DNA sites assuming closed packing on the surface (see S6).

**Figure 1.**
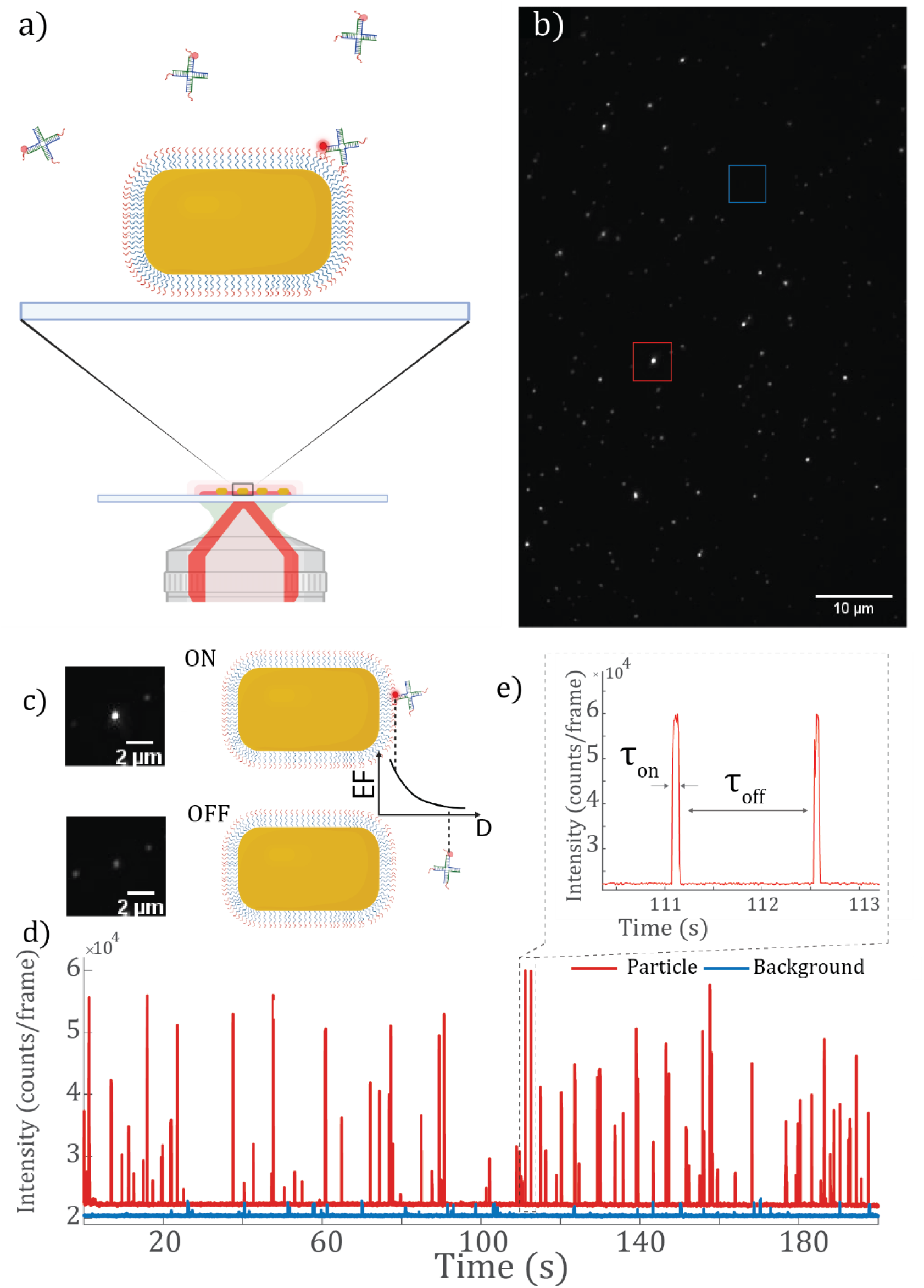
Measurement principle. a) Schematics of the setup. We use a DNA based framework of DNA coated Au nanorods and 4-way junctions (HJ) as a multivalent ligand-receptor system. The DNA coated nanorods are placed on a cover glass mounted in a flow cell on a TIRF microscope. When fluorescently labelled HJs bind to the particle through the DNA-DNA interaction the signal from a particle transiently increases. b) Example of a field of view showing the dark background and the diffraction limited bright spots from the gold nanorods. The blue square highlights a background region with HJ binding non-specifically shown in the blue time trace in d). c) Shows the nanorod highlighted in the red square in b) when a fluorescently labelled HJ is bound (top), referred to as the bound-state (on), and when no HJ is bound (bottom) which is the off-state. The sketch shows how the fluorescence enhancement factor (EF) drastically changes with the distance between particle and fluorophore-labeled HJ. d) shows a time trace from the highlighted particle in b) (red) and from the blue area with HJs binding to the background (blue). e) Zoom-in region of the time trace in d) that shows the bound-state lifetime τ_on_ (on-time/bright time) and off-time τ_off_ (dark-time) of the HJ binding to the nanorod

The gold particles appear as bright diffraction limited spots in the microscope due to one-photon luminescence (PL). The PL signal varies slightly from particle to particle due to differences in plasmon wavelength with respect to the laser excitation wavelength, among other effects, which results from inevitable polydispersity in wet-chemical synthesis protocols^37^. The PL signal makes a stable background and ensures each particle can be easily identified from the dark background. When a ligand DNA, carrying a fluorophore that spectrally overlaps with the plasmon, is bound to the gold nanorod its fluorescence gets enhanced and the diffraction spot brightness is greatly increased (figure 1C). We can thereby follow the signal from each particle over time and generate a time-trace from each particle as shown in figure 1d. The large intensity bursts come from single ligands binding to the particle. By tuning the concentration of molecules, we can make sure that each burst is from a single molecule. The intensity of the binding events varies which is expected as the molecules can bind to the particle at different locations where the field enhancement varies and hence the fluorescence enhancement differs.

The blue time trace in figure 1D comes from an ROI in the background i.e., not on a particle, where a fluorescent molecule (DNA ligand with fluorophore) binds nonspecifically to the surface and no plasmon enhancement of the fluorescence occurs. We can compare the intensity relative to the baseline to estimate the fluorescence enhancement. For the most intense events we observe an enhancement of up to almost 60 (see SI) which clearly highlights the advantage of using plasmon enhanced fluorescence to generate signals with superior brightness thereby enabling high signal-to-noise ratio even for short binding events.

Figure 1e shows a zoom-in of the time-trace. Following the methodology in qPAINT^29^ we can define the characteristic *bound-state lifetime* and *off-times* in the measurements. Binding events are identified when the intensity signal jumps above and below a threshold value. The *off-time(t_off_)*, also called the *dark-time*, is the time between binding events. The *bound-state lifetime* (*τ*_*on*_), also called *on-time*, is the time duration of a binding event. We focus on *τ*_*on*_where the avidity in multivalent binding is expected to greatly influence the duration of binding i.e., increase the binding strength.

### Multivalent kinetics from Holliday junctions

To study multivalency we engineered the number of ligand motifs in the interaction. We use DNA-DNA interactions as a model system for multivalent interactions, here addressing the case of a macromolecule binding to a surface that displays many receptors, e.g. MBL or C1q recognition of pathogen surfaces in the innate immune response^7^. Previous work has focused on kinetics of multivalent host-guest interactions of flexible macromolecules^23^ Here we deliberately work with a structured multivalent macromolecule to include the effects of the scaffold which is relevant as most biomacromolecules have a rigid structure. The model system has the benefit that we can tune the monovalent bound-state lifetimes (or equivalently the off-rate, *k*_*off*_) by the DNA sequence. We used a 4-way junction known as a Holliday Junction^38^ (HJ), specifically Holliday Junction 7^39,40^, as a scaffold. Nucleic acid scaffolds have gained increased interest^41^ because of their programmability, low cost and enhanced stability when introducing modified nucleic acids^42^. The HJ consists of four 22nt long strands that each bind to two other strands, schematically shown in figure 2a, that yields a 4-way junction. Following the nomenclature from the literature the arms are named R, B, H, and X^40^. Each arm can be modified to either carry the ligand sequence or a fluorescent label. Adjusting the length of the ligand sequence will change the binding strength. In this way we constructed mono-, bi-, and trivalent HJs (HJ1, HJ2, HJ3) and measured and compared the bound-state lifetimes of the individual constructs. We used two sets of sequences based on a commonly used sequence for DNA PAINT^28,30^ that are 8nt and 7nt long respectively. We used ATTO 655 as fluorescent tag on the 5’ end of HJ strand R. All ligand sequences were separated from the double stranded part of the HJ by a C nucleotide in order to avoid base stacking effects that greatly stabilize the interaction between the ligand and the receptor DNA ^43,44^.

**Figure 2.**
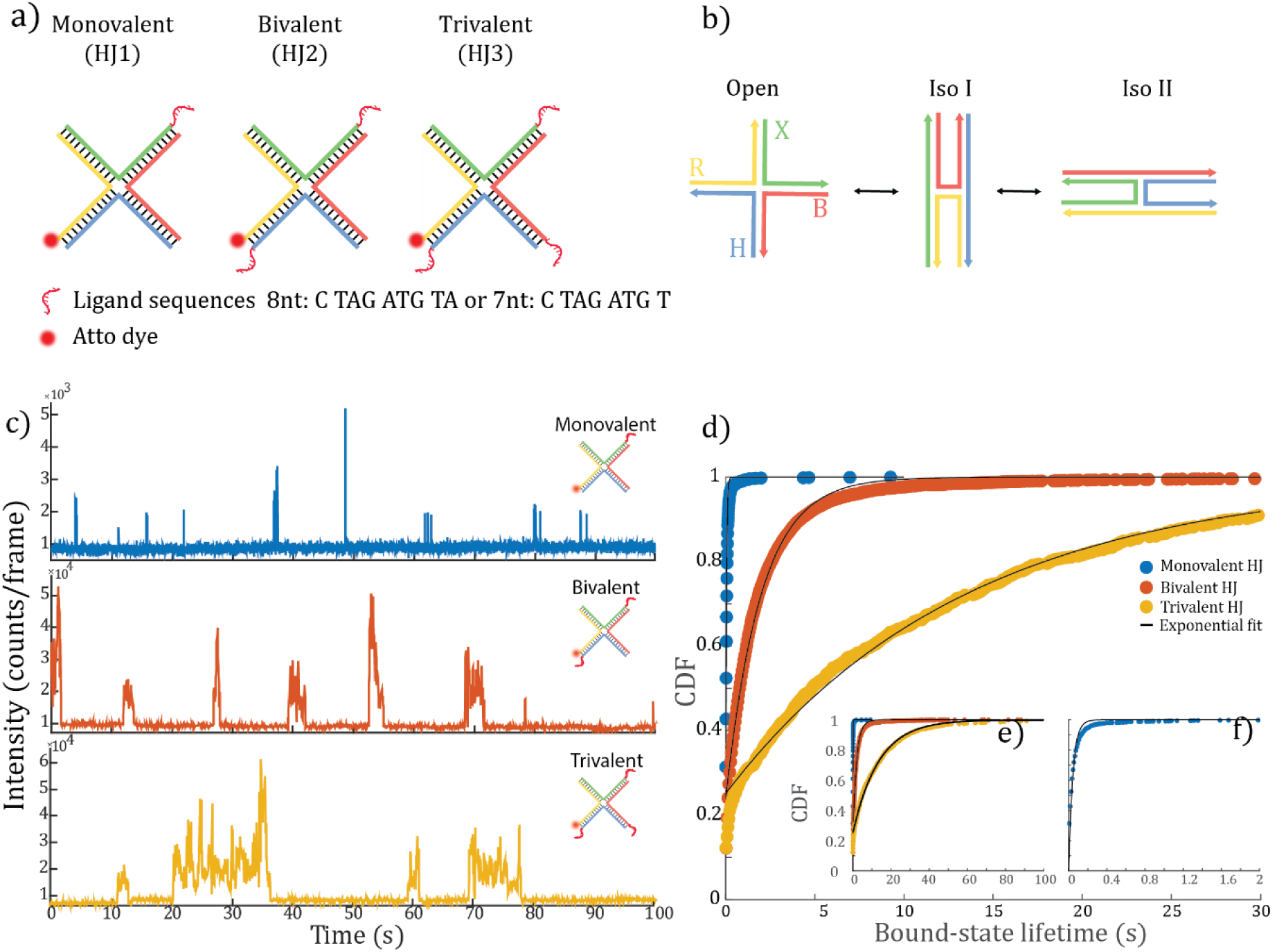
Holliday Junction with programmable valency. a) Sketch of the Holliday junction scaffold with dye location and added ligand sequences (ligands) showing the sequence that is complimentary to the receptor DNA on the gold nanorod surface. b) sketch of the open state and the two stacking isoforms iso I and II and the R, B, H and X arms. c) Examples of 100 second time traces from a monovalent (top), bivalent (middle) and trivalent (bottom) HJ with a 8nt ligand sequence. C) CDF of collected bound-state lifetimes from multiple time traces with thousands of events and a corresponding single exponential fit. e) inset of the CDF with longer time-axis to show the full trivalent curve. f) inset of the monovalent CDF on short time-scale.

Figure 2c shows exemplary time traces from a mono-, bi-, and trivalent HJ binding to the receptor DNA on the nanorods. From observing these times traces alone, it is clear that *τ*_*on*_increased drastically when the number of ligands was increased as expected for multivalent binding. For better comparison of bound-state lifetimes for the different HJs, we collected all events (thousands) from the single particles in each FOV and plot the cumulative distribution function (CDF) of all the lifetimes. We prepared samples such that there were around 50-100 particles in each FOV. Figure 2d shows the CDFs of a mono-, bi-, and trivalent HJ with an 8nt ligand. As was evident by the single time traces, the binding times drastically increased with the number of binding sequences, now taking into account thousands of binding events for each mono-, bi-, and trivalent HJ. The avidity effect from the multivalent binding is clear when considering the CDFs of the different constructs and will be subject to discussion later. To be sure this was in fact a multivalent effect we measured the binding of trivalent HJs to nanorods covered with a 1 to 50 ratio of receptor DNA and non-complementary DNA to physically space the receptor sites on the particle surface beyond the reach of the ligands on the HJs. In this case the CDF was similar to that of the monovalent HJ (see figure S11). It is therefore evident that the observed increase in bound-state lifetimes for the bi-and trivalent HJs in figure 2c-f was due to multivalent interactions.

To quantitatively compare the binding, we fitted the CDF for each construct with a single exponential function to extract the characteristic mean bound-state lifetime *τ*_*on*_using^37^: *CDF* = 1 − *A* ⋅ exp(*τ*/*τ*_*on*_) with A as a normalization factor. The off-rate *k*_*off*_ is directly linked to *τ*_*on*_ ^28^: *k*_*off*_ = ^1^. We can then compare the different constructs quantitatively via the extracted *τ* values from *τ on* the exponential fits. In the following sections we will look closer at the bivalent and trivalent HJs to get insights into the binding behavior.

### Bivalent interactions

First, we looked into theinteraction of the bivalent HJ (HJ2). Following the work of Huskens et al^23^, we describe the bivalent interaction of a single HJ as a step-wise binding mechanism as depicted in figure 3a. The first step is an intermolecular binding between one arm of the HJ and a receptor strand on the nanorod. The on-rate is determined by the on-rate of the single monovalent ligand (*k*_*on*,*M*_). After the first binding step the HJ can follow two routes. Firstly it can dissociate from the particle with an off-rate determined by the off-rate of the single ligand (*k*_*off,M*_). The second route is an intramolecular binding step with the second ligand where the on-rate depends on the effective concentration (*C*_*eff*_) ^18,45^ and the monovalent on-rate. The effective concentration is an important property in multivalent binding and can be thought of as the probability for the second ligand to find a receptor DNA that it can bind to. It is dependent on the length, L, of the bivalent object i.e., the distance between the two binding entities of a bivalent object and the surface density of receptor DNA. At macromolecular length scales the effective concentration is in the mM range. From the step-binding model we can describe the expected bound-state lifetime as (see Huskens, et. al^23^ and S7 for derivation):

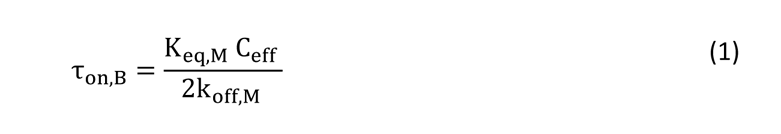

**Figure 3.**
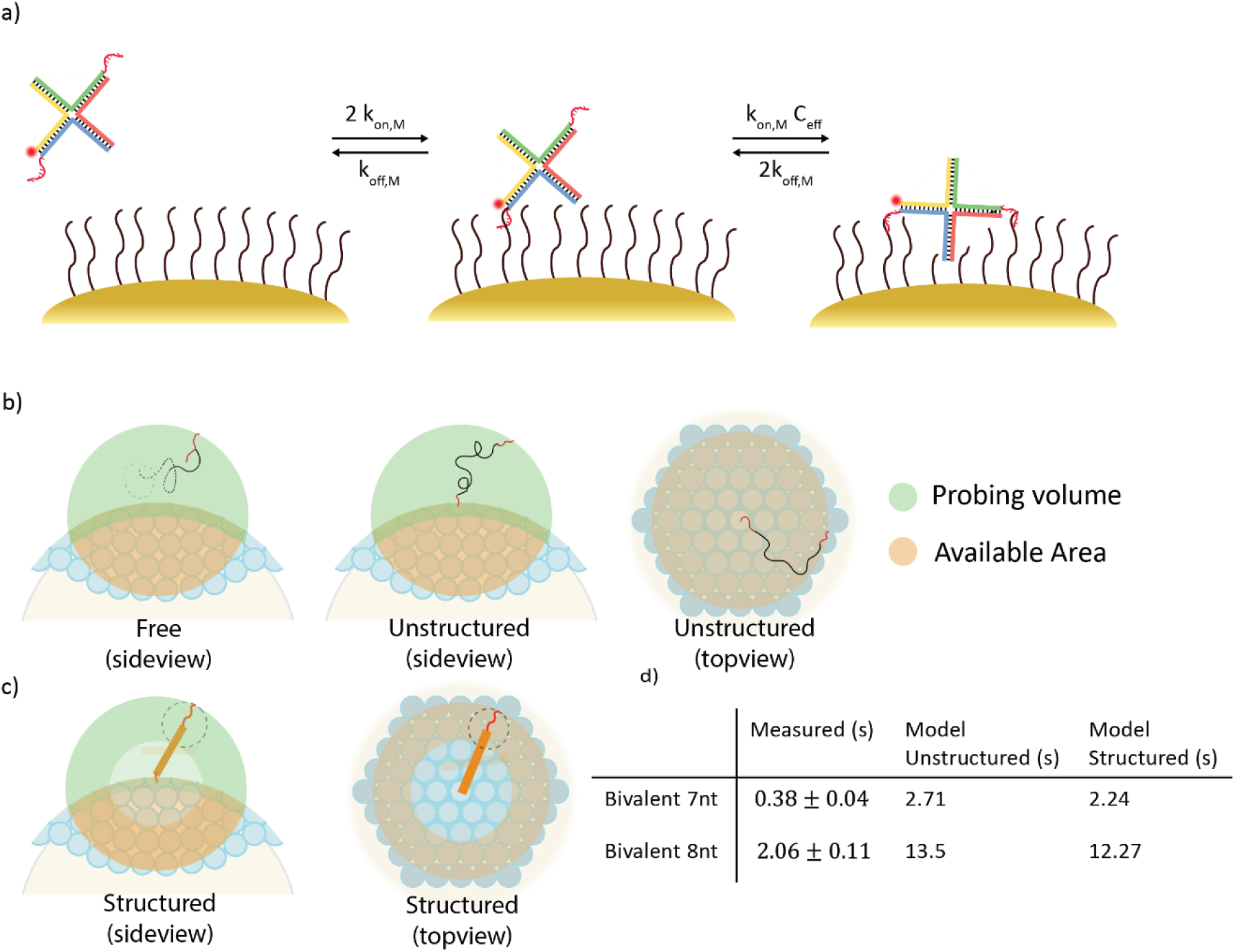
Step-binding model. a) Scheme of the step binding. First step is the binding of the HJ with one intermolecular interaction. From here the HJ can either dissociate and leave the particle or bind with the second ligand as an intramolecular interaction. The rate of the second binding step is influenced by the effective concentration (C_eff_). b) Schematics of the probing volume determining C_eff_. The green shows the probing volume and the orange shows the available surface area under the probing volume. The blue spheres (and half spheres) represent the receptor DNA on the particle surface. The first sketch (top left) shows the case when the ligand is considered a freely diffusing object within the probing volume. The second and third sketch shows the unstructured model based on the work of Huskens^23^ from the side and from the top. Here the ligand is tethered to the surface by the first ligand interaction but can diffuse freely within the probing volume defined by the length of the tether. c) Sketch of a structured model which takes into account the structure of the HJ that limits the accessible volume to be far from the first binding site. d) Experimental and modelled values of the bound-state lifetimes for bivalent HJs with 7nt and 8nt ligands.

Where *K*_*eq,M*_ is the monovalent binding constant. The model assumes independent interactions between ligandss and receptor DNA so that the individual interactions during the bivalent binding can be represented by the intrinsic on-or off-rates of the monovalent ligands. The predicted increasing bound-state lifetimes for the bivalent interactions are therefore due to high effective concentration and not because of cooperativity effects^46^.

We describe the effective concentration as the ratio between the probing volume of the ligand, V(L), and the number of available receptor sites on the surface, n_h_(L):

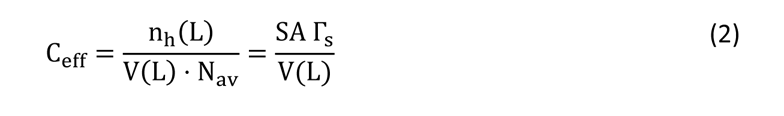

Where *N*_*av*_ is Avogadro’s number. The number of host sites can be expressed as the surface area of the particle-DNA surface, SA, and surface density of receptor DNA, Γ_s_. In our case we have to take into account the curved surface of the particle, which we do by considering the spherical ends of the particles where the plasmon enhancement is largely located (see figure 8c). To estimate the number of available receptor sites, *n*_ℎ_(*L*), on the reachable surface area, we used the measured average particle sizes and measured the DNA loading capacity on the particles using fluorescent receptor DNA quantified after removal from the particles^47^. A full description on the derivation of the effective concentration can be found in the supporting information (S7).

The probing volume is the available volume in which the molecule can move around. In the previous work from Huskens et al^23^, the probing volume was estimated by considering the ability of the second binding motif to move freely in a volume confined by the length of the polymer connecting the two ligandss (figure3b). It was presented as a tethered polymer but acting as a freely diffusing object where it is equally probable to reside at any place in the volume at any time. This model gave relatively good agreement to data for a flexible bivalent object^23^ relevant for intrinsically disordered proteins or unstructured polymer constructs, but likely cannot represent the structure present within most macromolecular proteins. We elaborated on the model to include structural considerations of the multivalent macromolecule. The HJ arms can stack on each other thereby giving rise to the two distinct isoforms I and II depicted in figure 2b. The HJ 7 used here has been reported to co-exist with almost equal isoform ratios at similar MgCl_2_ concentrations to those used here^40^. In our bivalent experiments the two ligand sequences are on the 3’ ends of X and H and therefore always far apart spanned by the length of the HJ. In a first attempt, following the previous work of Huskens et al, we used a probing volume within which the second ligand can move around freely. We have termed this the *unstructured* model. The span of the volume is determined by the length of the HJ and the ligand sequence. In the table in figure 3d are shown the measured bound-state lifetime values extracted from an exponential fit to the CDFs for the bivalent HJ with both the 7nt and 8nt binding sequences and the predicted values using the models. The increase in bound-state lifetimes from the monovalent to the bivalent HJ is around ∼ 30 and ∼ 40 times increase using 7nt and 8nt ligands respectively.

It is clear that the experimental values are much lower than the model values. Therefore, we further developed the model to consider that, due to the structure and rigidity of the HJ, there is a volume around the first binding which the second ligand cannot occupy and additionally a surface area (with corresponding number of receptors) that is not available for binding (figure 3c). This we refer to as the *structured* model. While the structured model does predict shorter bound-state lifetime values and so closer to the experimental values, they are a still a factor ∼ 6 longer than the experimental values.

### Restricted binding

Although the structured model, using simple geometrical considerations, represents data somewhat better, it does not yet describe how the macromolecular structure and conformation of the multivalent object affects the binding process. To better understand the multivalent binding behavior, we introduce a restriction factor *ω*. The concept of a restriction factor was introduced by Netz and Liese^19,48^ to describe the effect of ligand orientation when placed on the ends of polymeric chains. In our work the concept of the restriction factor is more general. Ultimately it accounts for all effects that restrict the binding capabilities of the second or subsequent binding interactions in multivalent binding when already bound through one or more interactions. We introduce the idea that instead of being equally probable at all sites in the probing volume, the second (third, …, etc.) ligand is likely to be in some parts of the probing space more often than in others. Figure 4a shows a conceptual sketch of the positional probability of the second ligand for the bivalent HJ and how the restriction factor affects the binding. Here, it is sketched such that the second ligand is less likely to be close to the surface, which is needed for it to bind, thus restricting its binding probability. We hypothesize that repulsion between the receptor DNA surface and the HJ makes it less likely to be at the surface, while steric hindrance within the HJ and stiffness of the HJ can restrict the availability of the ligand sequence. Therefore, the restriction factor is in general a property related to the structure of the multivalent object and to the binding interface that ultimately affects the effective concentration (*C*_*eff*_). Here we separate *C*_*eff*_ in two contributions; the first contribution is the binding restriction, as explained above, and the second contribution from being tethered through the already bound ligand, that defines a volume that the free ligand can span and the number of available receptors. The tethered contributions can thus be estimated as the *C*_*eff*_ from the unstructured model treating the ligand as a freely moving object within the tethered volume which we refer to as *C*^*free*^, thus estimated using eq. 2. With this separation we describe the effective concentration as *C*_*eff*_ = *C*^*free*^ ⋅ *ω*. Taking the expanded effective concentration using the binding restriction into account then yields a bound-state lifetime that depends on the restriction factor:

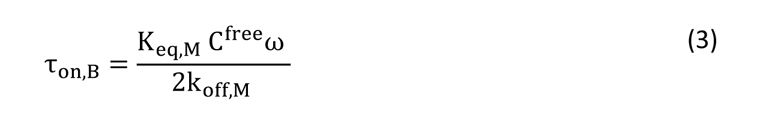

**Figure 4.**
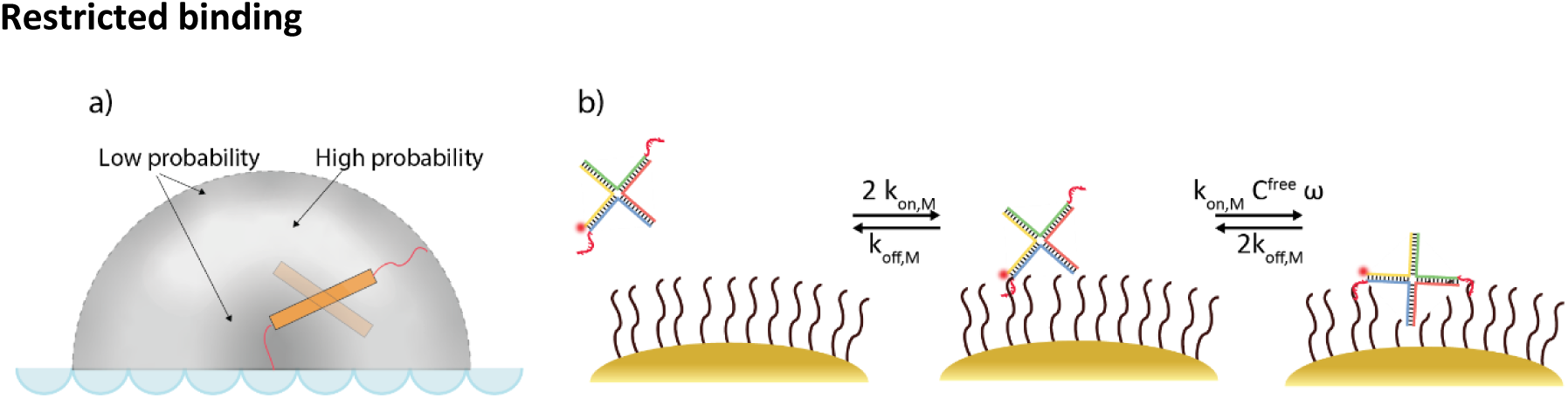
Restriction factor. a) Illustration of the influence of the restriction factor on the probability of the position of the second ligand during a binding event. The bright areas represent areas where the ligand is more likely to be, whereas the dark represents a volume the ligand is less probable to occupy. The restriction factor makes the space close to the surface less likely thus restricting the binding of the second ligand. b) Scheme of the step binding model with modified description of the effective concentration including the restriction factor, ω, and C^free^in the last step.

Note that a smaller value for the restriction factor means more restricted binding which gives rise to shorter bound-state lifetimes. While more complex models could be used this approach requires the least assumptions.

We can now use the model and our experiments to extract values for the restriction factor and allow comparison of the effect of changing the structure of the multivalent construct.

### Increasing ligand length and flexibility in the bivalent HJ

We wanted to apply our model to investigate how the length and flexibility of the binding arms affects the binding kinetics through restriction. To change the length, we introduced spacers into the sequence. The spacers consisted of different numbers of T nucleotides, from 2 to 18, between the ligand sequence and the HJ arm (figure 5a). For these measurements we used the 7nt sequence. The bound-state lifetimes and derived restriction factors for each construct are shown in figure 5b (left y-axis). There is a steep decrease in bound-state lifetime when adding the first T spacer. This is due to a decrease in *k*_*on*_ because of transient intrinsic hairpin formation that affects the *k*_*on*_^49^. We have used NUPACK^50^ to analyze the ligand sequences for self-interactions and found that when adding T spacers there is increased interaction between a C and G in the ligand sequences and that this C-G interaction is seemingly unchanged for all T-spacer lengths (see S4). To estimate how much *k*_*on*_ is reduced we have compared the event frequencies of 7nt ligands with and without T2 spacers. We do this by measuring from the same field of view (FOV) on our sample, i.e., the same particles. Using the event frequencies, we can then directly compare the *k*_*on*_ values and we have found that it drops 64% ± 16% when adding the T2 spacer. We used this to correct the *k*_*on*_ (*K*_*eq,M*_) values in eq.3 when adding spacers.

**Figure 5.**
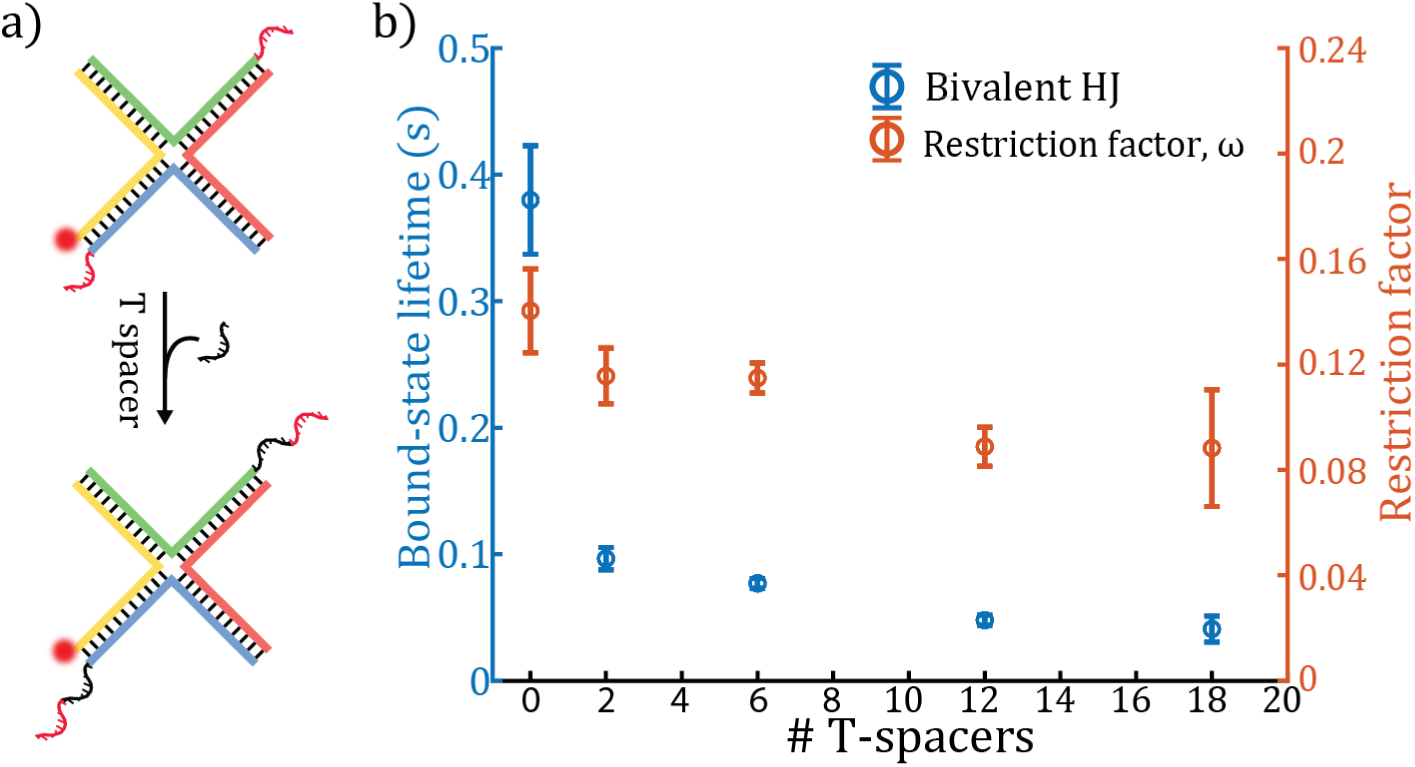
Addition of spacers. a) Spacer sequence addition between the HJ arm and ligand sequence that increases length and flexibility of the ligand. Spacer sequences consist of multiple T nucleotides ranging from 2 to 18. b) Plot of experimental bound-state lifetime values (left, blue) for the bivalent HJ with addition of different spacer lengths and the corresponding restriction factor (right, orange).

Further addition of T-spacers reveals a more steady decrease in bound-state lifetimes that approaches that of the monovalent HJ as the spacers become longer. This is explained by the expected decrease in effective concentration when increasing the spacer length^18^. In figure S9 are plotted the *C*^*free*^ changes with addition of T-spacer according to the *structured* and *unstructured* models. The graphs shows that addition of T-spacers reduces *C*^*free*^ due to an increased probing volume as expected^18^.The restriction factor gradually decreases with added spacer length which means binding of the second arm gets more restricted with longer spacer lengths. A possible explanation for the change in binding restriction is the orientation of the ligand that becomes less oriented towards the receptor surface. Note that Netz and Liese proposed that increased orientation of the ligand can increase the multivalent binding strength^19^. In their study they looked at systems with equal number of receptors and ligands in a matching geometry, which is not similar to our case in which we have a dense receptor surface. We hypothesize that the orientation of the ligand restricts its ability to reach the surface part close to where the HJ is bound. This effect then increases as more spacers are added.

### Trivalent binding

We then turned our attention to the trivalent binding HJ. Using the same approach for the bivalent HJ we describe the binding in a step-wise manner and now include an additional step with the third arm binding and corresponding restriction factors. The first two steps are essentially the same as in the bivalent case, except that the statistical pre-factors now take into account three available ligands initially. We here assume that during the first intramolecular transition, the second ligand binds far from the already bound ligand. For example, if the first binding interaction is established through the ligand on arm X, then we assume for the second step it binds through the ligand on arm H when in isoform I or either B or H when in isoform II. The *C*^*free*^ and the restriction factor for the second step established in the bivalent step-model can be used for those parameters in the trivalent step-model *C^free^_M→B_*.

A clear indication that the distant binding is preferred over the close binding in the second binding step is found by comparing two bivalent HJ constructs with ligands on the XH and XB arms. As described earlier the XH construct that we used for the bivalent measurements, will have ligandss distant in both isoforms, whereas for the XB construct the ligands are close in isoform I and far apart in isoform II. The two constructs show almost identical CDFs of the extracted bound-state lifetimes (see figure S11) which suggests that the XB preferably binds when in isoform II, i.e., when the ligands are distant.

When bound with two arms the HJ is brought in close contact with the surface. The third arm now experiences a *C*^*free*^ and binding restriction different from that in the second step. To distinguish between the restriction factors and *C*^*free*^in the two steps, the second step (bivalent binding) carries the subscript *M* → *B* (mono-to bivalent) and the third step *B* → *T* (bi-to trivalent). The derivation of the bound state time for the trivalent case is described in the supporting information (S7).

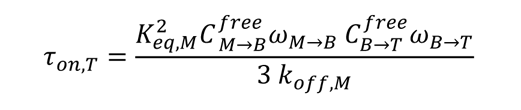

The probing volume for the third step is estimated using the length of the ligand sequence and, if added, any spacer length as well. This probe volume is significantly lower for the bivalent to trivalent transition compared to the monovalent to trivalent and the resultant C^free^ is significantly larger than the C^free^. The experimentally measured bound-state lifetimes for the trivalent HJs with 7nt and 8nt ligands are shown in the table in figure 6b. The restriction factors for the two multivalent steps are shown as well. The increase in lifetimes from the bivalent to the trivalent constructs is a factor ∼ 6 which is much less than what was observed from monovalent to bivalent binding (∼30-40). This relatively small bound-state lifetime increase is concurrent with a much more restricted binding for the third binding ligand(smaller restriction factor value). When already bound through two interactions the HJ is brought close to the surface and we hypothesize that it is therefore subject to steric effects on the third ligand arm that restrict its binding capabilities. Furthermore, we expect there to be an increased surface repulsion between the HJ and the DNA surface.

**Figure 6.**
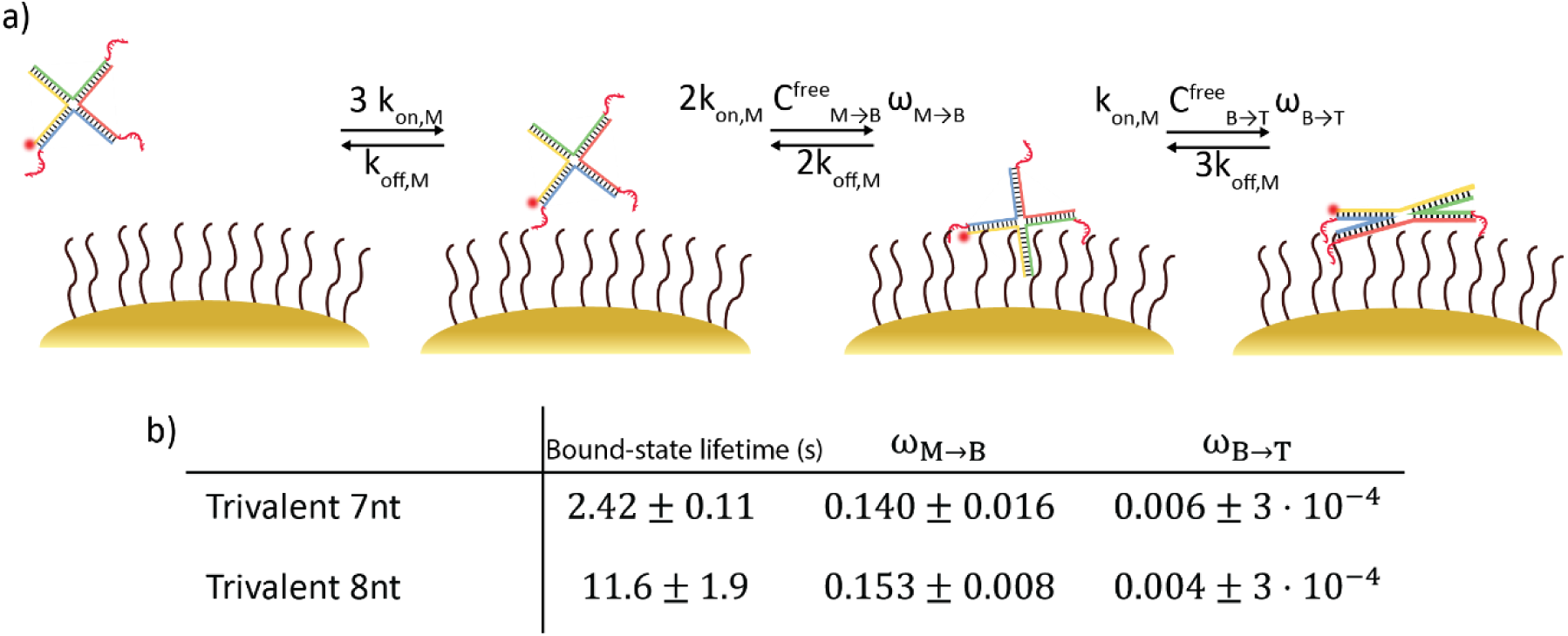
Trivalent step binding model. A) Scheme of the step binding model. First step is the binding of the HJ with one interaction. From there it can dissociate and leave the particle or bind with a second ligand under influence of the effective concentration C_eff_ = C^free^ ⋅ ω including the restriction factor ω associated with the monovalent to bivalent step (M → B). From this point it can either unbind with one of the interacting ligands or bind with the third ligand which is influenced by C^free^ and ω of the bivalent to trivalent step (B → T). B) Experimental bound-state lifetime values and restriction factor values for the trivalent HJ with 7nt and 8nt ligand sequences.

### Activating trivalent binding by decreasing binding restriction

Similarly to the bivalent HJ, we employ T-spacers on the trivalent HJ to explore the effect on binding restriction in the trivalent binding step. Figure 7b shows the bound-state lifetimes of the trivalent HJ with added spacers (left axis). Again, we observe a rapid drop when adding the first T-spacer, as for the bivalent HJ, which would be a consequence of a lowered *k*_*on*_ from the hairpin formation, which was explained for the bivalent case. The lifetimes decrease with added spacer length as expected due to the decrease in effective concentration (see S7) as was also observed for the bivalent HJ. Interestingly the decrease for the trivalent HJ with added spacer length is less pronounced compared to that for the bivalent HJ. We observe that the ratio of the bound-state lifetimes for the trivalent/bivalent HJs (see figure 7b in yellow) increases with spacer length and is doubled after addition of 6 T-spacer nucleotides. We interpret this to mean that the third binding arm on the trivalent HJs are very restricted and the additional T spacers can activate the binding, as seen by a reduction in the restriction. Figure 7c shows the binding restriction for the monovalent to bivalent (red) and bivalent to trivalent (black) binding steps. We have plotted the restriction as 1/*ω* to be more intuitive, such that a higher value refers to more binding restriction. Without any T-spacers the trivalent binding is very restricted compared to the bivalent. The restriction drops rapidly with the addition of T-spacers. It is worth noting that the changes in the bivalent restriction are weak compared to the trivalent. By adding the T-spacers we add flexibility that can release the steric effects present in the trivalent binding. Furthermore, the added T-spacers allow the HJ to be further away from the surface thus decreasing the surface repulsion. It should be noted that without the drop in *k*_*on*_ when adding T-spacers we would expect an even higher activation of the trivalent binding. Here we show that although addition of spacers leads in general to a decrease of bound-state lifetimes, the addition of spacers for trivalent HJs is more important as compared to bivalent as it can allow activation of the third binding arm. Addition of spacers for the bivalent HJs merely leads to a decrease of the effective concentration and lower bound-state lifetimes. We find this result interesting for understanding and designing multivalent ligands as previous work on the effects of linker lengths has, to our knowledge, mainly focused on the decrease in effective concentration and the role of rigidity and the resulting binding strengths^51,52^.

**Figure 7.**
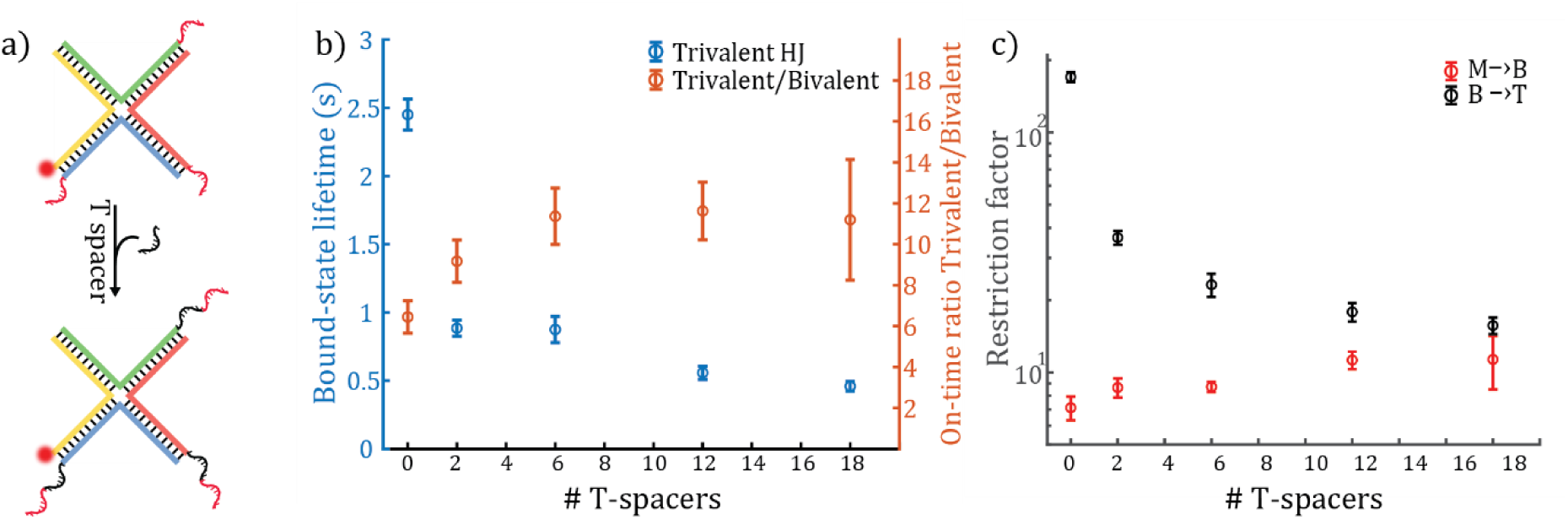
Addition of spacers in trivalent binding. A) Sketch of the added spacer sequences between the HJ arm and the ligand sequence. B) Experimental bound-state lifetime values for the trivalent HJ with added spacers (left, blue) and the bound-state lifetime value ratio between the trivalent and bivalent HJs with added spacers (right, orange). C) The evolution of restriction factor values, here plotted as 1/ω, for the monovalent to bivalent transition (red) and bivalent to trivalent (black) on a log-scale.

### Binding dynamics

So far, we have used the plasmon enhanced fluorescence signal only to detect binding events and quantify individual bound-state lifetimes. A powerful benefit, unique to a plasmon-enhanced approach, is the strong and rapidly decaying enhancement near the particle that induces a strong dependence of the fluorescence intensity on the exact location of the dye with respect to the particle. We exploit the strong distance-dependent signal of the plasmon enhanced fluorescence to give information about binding dynamics during the bound-state that may be encoded in the greatly fluctuating signal of the individual traces. This opens the potential to study the transitions between monovalent, bivalent and trivalent states to gain insights into the dynamic behavior of multivalent interactions.

During binding of the bivalent and trivalent HJs we observe signal fluctuations that are different from that of a monovalent binding signal (where no transitions during the bound state are expected). In figure 8a are shown exemplary time traces from a 12nt monovalent ligand and in figure 8b the trivalent HJ with 8nt ligands. The signal from the monovalent ligand is most often a box-like signal which, when binding occurs, reaches one level that it fluctuates around. The difference in fluorescence signal between individual binding events is expected as the monovalent ligand can bind at different locations on the metal particle where different field strengths are expected, giving differences in fluorescence enhancement. In contrast, the signal during a single binding event of the trivalent HJ most often does not remain at a single fluorescence intensity level but shows many jumps between fluorescence levels during the binding event before finally unbinding. The fluctuations from the 12nt ligand (figure 8a) are most likely due to dye blinking, resulting in a signal that digitally fluctuates between zero and non-zero values, which is expected to be present at a similar level in the trivalent HJ signal (figure 8b). However, the monovalent and trivalent traces are very different which can be seen from the histograms of the intensity levels in figure 8a-b. The trivalent HJ events are spread out over a larger range of fluorescence intensity levels compared to the 12nt monovalent ligand, which is evident by the histogram and the corresponding relative standard deviations (RSD). The plasmon enhanced signal depends on the position of the dye relative to the particle^33,34^ which can clearly be seen from the BEM simulation in figure 8c that shows the plasmon enhancement for a single 40 nm by by 82 nm nanorod. The strongest signal is reached around the ends of the nanorod where it can reach fluorescence enhancement factors above 70. The cross-sectional plot of the enhancement factors along the black line shows that the enhancement factor changes greatly depending on the position along the line and the maximum value is reached around a 3.5nm distance from the particle (figure 8d). In our case the fluorophore is likely to be no closer than 5 nm due to the receptor DNA layer and the HJ scaffold. Taking this into account the observed fluorescence enhancement factors are in good agreement with the simulated values.

**Figure 8.**
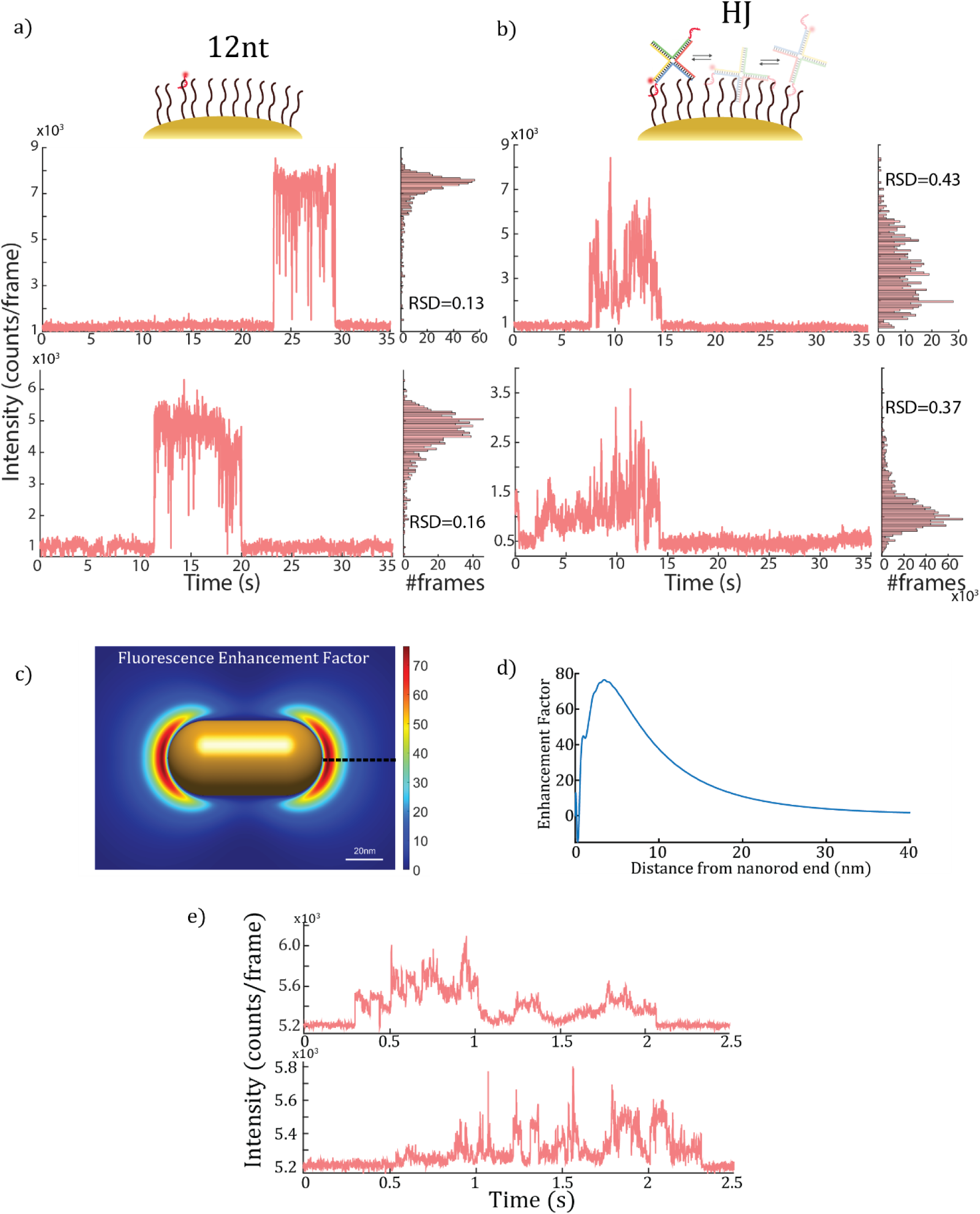
Binding dynamics. a) Time traces of a 12nt monovalent ligand showing a ‘box-like signal’ with fluctuations around a single fluorescence intensity level within an event. On the right is plotted a histogram of the signal intensities for each frame along with the relative standard deviation (RSD) of the signal values. b) Time traces of a trivalent HJ with 8nt ligands that show different signal fluctuations compared to the 12nt ligand. Here the signal fluctuates around multiple fluorescence intensity levels. On the right is plotted a histogram of the signal intensities for each frame along with the relative standard deviation (RSD) of the signal values. Traces in a) and b) were acquired with 100 fps framerate. c) BEM simulation of a 40 by 82 nm Au nanorod with LSPR at 650nm showing the fluorescence enhancement factor of an ATTO 655 dye with an intrinsic quantum yield of 0.3. d) Fluorescence enhancement factor along the dotted line in c) that highlights the distance dependent enhancement. e) Zoom-in of trivalent HJ binding events with 1000 fps from a cropped FOV.

We propose that the fluctuations observed in the trivalent binding signal are due to the changing position of the dye during the dynamic stepwise binding. When the trivalent HJ is bound, each arm of the HJ can bind and unbind multiple times that should change the position of the dye in each step. The time scale of the transitions between mono-, bi-, and trivalent binding during a single binding event can be estimated from the step binding model. The estimated values for the trivalent HJ with 8nt ligands are listed in the S8 in the supporting information. The transition rates for the unbinding steps are 2 or 3 times faster than the monovalent due to the additional statistical factors. Because of the much faster transition from mono-to bivalent binding, we expect the HJ to be bound in the trivalent state most of the time within a binding event and transition between the trivalent and bivalent states. Using a 10ms framerate, which is the fastest possible in the setup that we have used for all time traces shown thus far, we expect on average a binding step event in each frame. Due to the distance dependent signal enhancement, the signal will change for every transition and so we would expect drastically different signal fluctuations for the trivalent HJ compared to the 12nt imager. It has been proposed that multivalent objects can walk and even hop around on the surface^53^ which would also give rise to signal fluctuations, likely on a slower time scale as the HJ gradually explores areas of varying enhancement. Here we have shown proof of principle of the measurement of dynamics during binding events of multivalent ligands.

Using a custom-built TIRF microscope with a cropped FOV on the CMOS camera, we were able to acquire time traces with 1000 fps of a trivalent HJ shown in figure 8e (more events are shown in S16 in the supporting information). These events show that even with such a high acquisition rate, binding events are easily detected from the baseline signal. Furthermore, looking at the time traces we can clearly see steps and plateaus of the intensity signal on timescales around 5-50ms likely showing the waiting time between a transition from one bound state to another. The time resolution of fluorescence measurements can in principle be improved in particular exploiting the relatively high plasmon enhanced signals to reach timescales relevant for studying individual transitions in bound states during multivalent binding events.

## Conclusion

In this work we have used plasmon enhanced fluorescence to study the kinetics and dynamics of multivalent interactions on the single-molecule level. Using a model system formed from a DNA scaffold (Holliday Junction), we have applied the plasmon enhanced approach to study monovalent, bivalent and trivalent interactions and quantitatively extract bound state lifetimes. We have developed a model framework and introduced the general concept of binding restriction to take into account how the structure and molecular conformation of multivalent ligands affect the effective concentration and the binding. For our system we observe that the bivalent to trivalent transition is subject to a great amount of binding restriction. By comparing the effect of increased linker length and flexibility on bivalent and trivalent constructs we confirmed the general effect that the addition of spacers leads to reduced bound state lifetimes through a resultant decrease in the effective concentration of the binding motif. Interestingly we found that trivalent binding was activated and much less restricted with adding length and flexibility to the binding arms giving the potential for increased selectivity. Furthermore, we demonstrated the measurement of binding dynamics within single binding events of multivalent systems by utilizing the distance dependent plasmon enhanced fluorescence to probe changes in the dye position that arise from the binding dynamics. This approach has the potential for broad application to provide quantitative insight into multivalent interactions and guide the design of multivalent synthetic macromolecules for use in therapeutic or bioimaging applications.

## Authors

**Kasper R. Okholm** - Interdisciplinary Nanoscience Center, Aarhus University, Gustav Wieds Vej 14, Aarhus C, 8000, Denmark. The Centre for Cellular Signal Patterns (CELLPAT), Gustav Wieds Vej 14, Aarhus C, 8000, Denmark

**Sjoerd W. Nooteboom** - Eindhoven University of Technology, Department of Applied Physics and Science Education and Institute for Complex Molecular Systems, 5600 MB Eindhoven, The Netherlands

**Vincenzo Lamberti** - Eindhoven University of Technology, Department of Applied Physics and Science Education and Institute for Complex Molecular Systems, 5600 MB Eindhoven, The Netherlands

**Swayandipta Dey** - Eindhoven University of Technology, Department of Applied Physics and Science Education and Institute for Complex Molecular Systems, 5600 MB Eindhoven, The Netherlands

## Author contributions

K.R.O and S.W.N contributed equally to this work.

## Acknowledgements

This work acknowledges funding from the Danish National Research Foundation center grant CellPAT (DNRF135) (K.R.O, D.S.S). This project has received funding from the European Research Council (ERC) under the European Union’s Horizon 2020 research and innovation programme (grant agreement No 864772). V.L. and P.Z. acknowledge funding from the European Union’s Horizon 2020 research and innovation program under the Marie Skłodowska-Curie program (ITN SuperCol, Grant Agreement 860914).

## Materials and methods

### Materials

DNA strands labelled with ATTO655 and ATTO643 were purchased from Eurofins genomics. All other DNA strands were purchased from Integrated DNA Technologies (IDT). TAE was purchased from Gibco and both MgCl_2_ and NaCl were purchased form Sigma Aldrich. BSA-biotin and Neutravidin were purchased from Thermo scientific. Gold nanorods (A12-40-650 citrate) were purchased from Nanopartz. Accugel 40% was purchased from National Diagnostics and we used Ultra-low range ladder from Invitrogen. Gold etchant was purchased from Sigma Aldrich.

### Optical setup

We used an Oxford nanoimager (ONI) microscope operating in TIRF mode equipped with a 100x/1.4NA oil immersion objective. The microscope was gradually heating when starting imaging at room temperature. Therefore, we performed all imaging at 27 °C at which the temperature was stable throughout the imaging sessions. For all experiments, except the 1000 fps acquisition, we used a 640 nm laser as excitation source at 20 mW power (*power densisty:* 320*w/m^2^*). Each image was recorded on a 427x520-pixel region with 117 nm pixel size.

For the 1000 fps data we used an objective-type total internal reflection microscope on a Nikon Ti2 microscope with an oil-immersion objective (Nikon Apo TIRF 60x Oil DIC N2, 1.49 NA). A 637 nm excitation laser (OBIS FP 637LX, Coherent) was used to excite the sample in s polarization via a dichroic mirror (ZT640rdc, Chroma). The maximum power density on the sample was approximately 2 ⋅ 10^6^ *w/m^2^* to avoid the influence of photothermal heating on the dynamics. Fluorescence was collected through the same objective and the leftover laser reflection was removed by a notch filter (ZET635NF, Chroma) and a long-pass filter (FELH0650, Thorlabs). The signal was detected on a CMOS camera (Teledyne Photometrics Prime BSI Express), in 11-bit mode and cropped to 128x128 pixels to allow for 1000 frames per second acquisition. Note that for the 1000 fps acquisition we have used ATTO643 labeled HJs.

### Data analysis

To generate time traces we used in-house software as describes previously^30,37^. In brief, the fluorescence time traces were generated by summing the intensity in a 7x7 pixel region from identified particles in each frame. Time trace analysis was carried out using home-made MATLAB scripts. We employed a threshold value and frames above the threshold were identified as events and the duration as the number of consecutive frames within the event multiplied by the camera acquisition time. As each event is counted in integer number of frames, we correct the duration of the last frame to only last half the acquisition time to avoid overestimating the duration of fast events. The CDF of bound-state lifetimes were fitted to *CDF* = 1 − *Ae*^−*τ*/*τ on*^, where *τ*_*on*_is the mean bound-state lifetime. For the fast events (7nt and 8nt imagers and monovalent HJs) where lifetimes were close to the camera frame duration, we used A=1 and excluded the first datapoint in the CDF (which is poorly defined and makes up many of the total events) from the fit. For each construct we measured in three different FOV and therefore get three CDFs and extracted lifetime values. The reported bound-state lifetime values are the average of the three lifetime values and the error is the standard deviation of the three lifetime values for each construct.

### Holliday junction assembly

For the Holliday junction assembly we mixed the four strands (R, B, H and X) at equimolar concentrations in 1xPBS and then heated to 70°C followed by a linear temperature cooling to 4°C over 90 minutes using a thermocycler. All assembled constructs were analyzed by 12% Native PAGE using 200V for 2h with 1xTBE as running buffer and appr. 0.5 pmol amount of HJ construct. The gel was stained and imaged with an Amersham Typhoon laser scanner.

### Gold particle DNA coating preparation

We have used a freeze-thawing approach to coat the nanorods with DNA^36^. First we mixed TCEP and Thiol-DNA for 5 min with 200X excess TCEP. The DNA and TCEP was added to an eppendorf tube containing the citrate capped AuNR suspension. The DNA was appr. 2.5⋅ 10^5^X excess to AuNRs. The suspension was briefly mixed on a vortex and then we added SDS to a final concentration of 0.01%SDS. We vortexed and centrifuged briefly and put the suspension on dry-ice for few minutes until frozen. After freezing the suspension was left at room-temperature to thaw. Then we repeated the freeze-thawing cycles once more. Here, we kept the particle suspension on freeze until approximately an hour before use. After thawing we centrifuged the particles at 10.000x G for 2 minutes such that the particles formed an oily pellet. We carefully removed the supernatant and resuspended in 1xTAE 0.01%SDS. We repeated the centrifugation and resuspension 3 times.

### Sample preparation

We have used 24x60 mm #1.5 microscope cover glasses as sample substrate. First, we flushed the slides with acetone and then ethanol for 10 seconds each. After that we blow dried the slides and further cleaned in a UV/Ozone oven for approximately 1h. Then we attached ibidi sticky slides (sticky slide VI 0.4, ibidi) on the clean glass surface to make our flowcells. We functionalized the surface with BSA-biotin, Neutravidin and receptor DNA strands, by sequentially adding the different solutions to the channel and washing in between. First we added 50 μL of 1mg/mL BSA-biotin in 1xTAE with 100mM NaCl to the channel and incubated for 5 minutes. The channel was then washed with 100 μL 1xTAE with 100mM NaCl twice. Then we added 50 μL 0.5 mg/mL in 1xTAE with 100mM NaCl Neutravidin to the channel, incubated for 3 minutes and again followed by a double washing step. After this we added 50 μL of 25nM receptor DNA in 1xTAE with 100mM NaCl and washed four times after incubating for 3 minutes. Then we added the gold particle suspension to the channel. The particle coverage is tuned by the concentration of the particle suspension and incubation time. Usually, we would dilute the particle suspension to half of the concentration obtained right after the purification step after freeze-thawing. We would add 30 uL of the suspension to the channel and wash twice with 1xTAE with 10mM MgCl_2_after 10 sec. incubation.

### Imaging conditions

Depending on the bound-state lifetimes of the HJ or single ligand we would use 10pM-2nM in 1xTAE with 10mM MgCl_2_ to be sure we were observing single events. For constructs with short bound-state lifetimes we used high sample concentration and 10ms camera integration time (100*S*^−1^framerate) whereas for constructs with long bound-state lifetimes we used low concentration and 50ms camera integration time. We recorded 10000-20000 frames for each FOV depending on the event frequency (which depends on the concentration). All measurements that are used in comparison were conducted the same or consecutive to have as identical conditions as possible for the measurements in comparison. We used ATTO655 labeled HJs for all experiments except the measurements using 1000fps where we have used ATTO643 instead.

### Simulations

We carried out numerical simulation of the optical proprieties of NP and fluorescence enhancement using the boundary element method (BEM), with the MNPBEM17 toolbox^54^, as previously described^37^. Here we used a 40 by 82 nm nanorod that was modeled as a cylinder capped by hemispheres and the Atto655 fluorophore with 30% quantum yield was modeled as a point-like dipole where three perpendicular orientations of the dipole were considered and averaged in resulting quantities, assuming that the dye rotates faster than its fluorescence lifetime. Light interaction of the NR and fluorophore were calculated with 637 nm illumination polarized along the long axis of the rods. A total of 82x40 relative positions between NP and dipole were simulated. We applied a 2D interpolation function to obtain high-resolution images of total fluorescence enhancement. Decay rate enhancements were calculated over the entire emission spectrum of the dye. The total fluorescence enhancement was calculated by multiplying the near-field intensity enhancement and quantum yield modification, neglecting fluorescence saturation.

## Bibliography

1. Mammen, M., Choi, S.-K. & Whitesides, G. M. Polyvalent Interactions in Biological Systems: Implications for Design and Use of Multivalent Ligands and Inhibitors. Angew. Chem. Int. Ed. 37, 2754–2794 (1998).

2. Fasting, C., et al. Multivalency as a Chemical Organization and Action Principle. Angew. Chem. Int. Ed. 51, 10472–10498 (2012).

3. *Multivalency: Concepts, Research & Applications*. (Wiley, 2018). doi:10.1002/9781119143505.

4. Haag, R. Multivalency as a chemical organization and action principle. Beilstein J. Org. Chem. 11, 848–849 (2015).

5. Speziale, P. & Pietrocola, G. The Multivalent Role of Fibronectin-Binding Proteins A and B (FnBPA and FnBPB) of Staphylococcus aureus in Host Infections. Front. Microbiol. 11, (2020).

6. Stencel-Baerenwald, J. E., Reiss, K., Reiter, D. M., Stehle, T. & Dermody, T. S. The sweet spot: defining virus–sialic acid interactions. Nat. Rev. Microbiol. 12, 739–749 (2014).

7. Bajic, G., Degn, S. E., Thiel, S. & Andersen, G. R. Complement activation, regulation, and molecular basis for complement-related diseases. EMBO J. 34, 2735–2757 (2015).

8. Satav, T., Huskens, J. & Jonkheijm, P. Effects of Variations in Ligand Density on Cell Signaling. Small 11, 5184–5199 (2015).

9. Woythe, L., Tito, N. B. & Albertazzi, L. A quantitative view on multivalent nanomedicine targeting. Adv. Drug Deliv. Rev. 169, 1–21 (2021).

10. Tjandra, K. C. & Thordarson, P. Multivalency in Drug Delivery–When Is It Too Much of a Good Thing? Bioconjug. Chem. 30, 503–514 (2019).

11. Krishnamurthy, V. M., Estroff, L. A. & Whitesides, G. M. Multivalency in Ligand Design. in (2006). doi:10.1002/3527608761.ch2.

12. Böhmer, V. I., Szymanski, W., Feringa, B. L. & Elsinga, P. H. Multivalent Probes in Molecular Imaging: Reality or Future? Trends Mol. Med. 27, 379–393 (2021).

13. Badjić, J. D., Nelson, A., Cantrill, S. J., Turnbull, W. B. & Stoddart, J. F. Multivalency and Cooperativity in Supramolecular Chemistry. Acc. Chem. Res. 38, 723–732 (2005).

14. Xiu, F. et al. Multivalent Noncovalent Interfacing and Cross-Linking of Supramolecular Tubes. Adv. Mater. 34, 2105926 (2022).

15. Khateb, H. et al. The Role of Nanoscale Distribution of Fibronectin in the Adhesion of *Staphylococcus aureus* Studied by Protein Patterning and DNA-PAINT. ACS Nano 16, 10392–10403 (2022).

16. Kitov, P. I. & Bundle, D. R. On the Nature of the Multivalency Effect: A Thermodynamic Model. J. Am. Chem. Soc. 125, 16271–16284 (2003).

17. Kiessling, L. L., Gestwicki, J. E. & Strong, L. E. Synthetic multivalent ligands in the exploration of cell-surface interactions. Curr. Opin. Chem. Biol. 4, 696–703 (2000).

18. Huskens, J. et al. A Model for Describing the Thermodynamics of Multivalent Host−Guest Interactions at Interfaces. J. Am. Chem. Soc. 126, 6784–6797 (2004).

19. Liese, S. & Netz, R. R. Quantitative Prediction of Multivalent Ligand–Receptor Binding Affinities for Influenza, Cholera, and Anthrax Inhibition. ACS Nano 12, 4140–4147 (2018).

20. Milroy, L.-G., Grossmann, T. N., Hennig, S., Brunsveld, L. & Ottmann, C. Modulators of Protein–Protein Interactions. Chem. Rev. 114, 4695–4748 (2014).

21. H. (Erik) Hamming, P., J. Overeem, N. & Huskens, J. Influenza as a molecular walker. Chem. Sci. 11, 27–36 (2020).

22. Riera, R. et al. Single-molecule imaging of glycan–lectin interactions on cells with Glyco-PAINT. Nat. Chem. Biol. 17, 1281–1288 (2021).

23. Gomez-Casado, A. et al. Probing Multivalent Interactions in a Synthetic Host–Guest Complex by Dynamic Force Spectroscopy. J. Am. Chem. Soc. 133, 10849–10857 (2011).

24. Lallemang, M. et al. Multivalent non-covalent interactions lead to strongest polymer adhesion. Nanoscale 14, 3768–3776 (2022).

25. Howorka, S., Nam, J., Bayley, H. & Kahne, D. Stochastic Detection of Monovalent and Bivalent Protein–Ligand Interactions. Angew. Chem. Int. Ed. 43, 842–846 (2004).

26. Semeniak, D., Cruz, D. F., Chilkoti, A. & Mikkelsen, M. H. Plasmonic Fluorescence Enhancement in Diagnostics for Clinical Tests at Point-of-Care: A Review of Recent Technologies. Adv. Mater. 2107986 (2022) doi:10.1002/adma.202107986.

27. Wang, Y., Horacek, M. & Zijlstra, P. Strong Plasmon-Enhancement of the Saturation Photon Count Rate of Single Molecules. J. Phys. Chem. Lett. (2020).

28. Jungmann, R. et al. Single-Molecule Kinetics and Super-Resolution Microscopy by Fluorescence Imaging of Transient Binding on DNA Origami. Nano Lett. 10, 4756–4761 (2010).

29. Jungmann, R. et al. Quantitative super-resolution imaging with qPAINT. Nat. Methods 13, 439–442 (2016).

30. Horáček, M., J. Engels, D. & Zijlstra, P. Dynamic single-molecule counting for the quantification and optimization of nanoparticle functionalization protocols. Nanoscale 12, 4128– 4136 (2020).

31. Horáček, M., Armstrong, R. E. & Zijlstra, P. Heterogeneous Kinetics in the Functionalization of Single Plasmonic Nanoparticles. Langmuir 34, 131–138 (2018).

32. Maier, S. A. Plasmonics: Fundamentals and Applications. (Springer US, 2007). doi:10.1007/0-387-37825-1.

33. Anger, P., Bharadwaj, P. & Novotny, L. Enhancement and Quenching of Single-Molecule Fluorescence. Phys. Rev. Lett. 96, 113002 (2006).

34. Kühn, S., Håkanson, U., Rogobete, L. & Sandoghdar, V. Enhancement of Single-Molecule Fluorescence Using a Gold Nanoparticle as an Optical Nanoantenna. Phys. Rev. Lett. 97, 017402 (2006).

35. Khatua, S. et al. Resonant Plasmonic Enhancement of Single-Molecule Fluorescence by Individual Gold Nanorods. ACS Nano 8, 4440–4449 (2014).

36. Liu, B. & Liu, J. Freezing Directed Construction of Bio/Nano Interfaces: Reagentless Conjugation, Denser Spherical Nucleic Acids, and Better Nanoflares. J. Am. Chem. Soc. 139, 9471– 9474 (2017).

37. Nooteboom, S. W., Wang, Y., Dey, S. & Zijlstra, P. Real-Time Interfacial Nanothermometry Using DNA-PAINT Microscopy. Small 18, 2201602 (2022).

38. Holliday, R. A mechanism for gene conversion in fungi. Genet. Res. 5, 282–304 (1964).

39. Grainger, R. J., Murchie, A. I. H. & Lilley, D. M. J. Exchange between Stacking Conformers in a Four-Way DNA Junction. Biochemistry 37, 23–32 (1998).

40. McKinney, S. A., Déclais, A.-C., Lilley, D. M. J. & Ha, T. Structural dynamics of individual Holliday junctions. Nat. Struct. Biol. 10, 93–97 (2003).

41. B. Yeldell, S. & Seitz, O. Nucleic acid constructs for the interrogation of multivalent protein interactions. Chem. Soc. Rev. 49, 6848–6865 (2020).

42. Omer, M., Andersen, V. L., Nielsen, J. S., Wengel, J. & Kjems, J. Improved Cancer Targeting by Multimerizing Aptamers on Nanoscaffolds. Mol. Ther. - Nucleic Acids 22, 994–1003 (2020).

43. Banerjee, A., Anand, M., Kalita, S. & Ganji, M. Single-Molecule Analysis of DNA Base-Stacking Energetics Using Patterned DNA Nanostructures. Preprint at 10.1101/2022.09.08.506950 (2022).

44. Abraham Punnoose, J., et al. High-throughput single-molecule quantification of individual base stacking energies in nucleic acids. Nat. Commun. 14, 631 (2023).

45. Krishnamurthy, V. M., Semetey, V., Bracher, P. J., Shen, N. & Whitesides, G. M. Dependence of Effective Molarity on Linker Length for an Intramolecular Protein−Ligand System. J. Am. Chem. Soc. 129, 1312–1320 (2007).

46. Ercolani, G. Assessment of Cooperativity in Self-Assembly. J. Am. Chem. Soc. 125, 16097–16103 (2003).

47. Demers, L. M. et al. A Fluorescence-Based Method for Determining the Surface Coverage and Hybridization Efficiency of Thiol-Capped Oligonucleotides Bound to Gold Thin Films and Nanoparticles. Anal. Chem. 72, 5535–5541 (2000).

48. Lauster, D. et al. Phage capsid nanoparticles with defined ligand arrangement block influenza virus entry. Nat. Nanotechnol. 15, 373–379 (2020).

49. Schueder, F. et al. An order of magnitude faster DNA-PAINT imaging by optimized sequence design and buffer conditions. Nat. Methods 16, 1101–1104 (2019).

50. Zadeh, J. N. et al. NUPACK: Analysis and design of nucleic acid systems. J. Comput. Chem. 32, 170–173 (2011).

51. Bandlow, V. et al. Spatial Screening of Hemagglutinin on Influenza A Virus Particles: Sialyl-LacNAc Displays on DNA and PEG Scaffolds Reveal the Requirements for Bivalency Enhanced Interactions with Weak Monovalent Binders. J. Am. Chem. Soc. 139, 16389–16397 (2017).

52. Matsui, M. & Ebara, Y. Enhanced binding of trigonal DNA–carbohydrate conjugates to lectin. Bioorg. Med. Chem. Lett. 22, 6139–6143 (2012).

53. Perl, A. et al. Gradient-driven motion of multivalent ligand molecules along a surface functionalized with multiple receptors. Nat. Chem. 3, 317–322 (2011).

54. Hohenester, U. & Trügler, A. MNPBEM – A Matlab toolbox for the simulation of plasmonic nanoparticles. Comput. Phys. Commun. 183, 370–381 (2012).

