## Supporting Information for "Kinetics and dynamics of single-molecule multivalent interactions revealed by plasmon-enhanced fluorescence"

#### S1. Holliday Junction

The Holliday junction used here is referred to in the literature as Junction 7[1,2]. Each sequence is termed X, B, R and H as shown in Figure S1. In the presence of salt, the HJ adopts two distinct isoforms in which the arms can stack in different ways and can transition between these two isoforms. It has been shown that the Junction 7 co-exist in the two isoforms at almost equal ratios[2] with a transition rate of about $10s^{-1}$ under similar buffer conditions as used in this work.

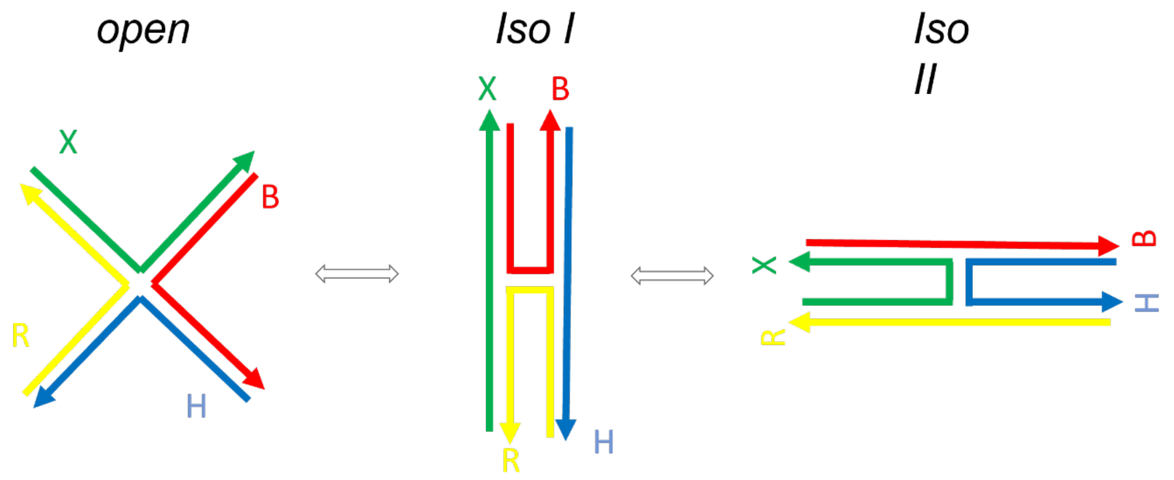

*Figure S1 HJ stacking isoforms. The figure sketches the different isoforms of the Holliday Junction. By introduction of salt in the HJ solution, the arms can stack and it transitions from the open state between the two isoforms I and II (Iso I and Iso II).*

List of sequences:

*Table S1 List of DNA sequences used*

| Strand name | Sequence (5' -> 3') |
|---|---|
| R | CCCACCGCTCGGCTCAACTGGG |
| R-atto655 5' | 5'-atto 655- CCCACCGCTCGGCTCAACTGGG-'3 |
| X | CCCAGTTGAGCGCTTGCTAGGG |
| X 8nt | CCCAGTTGAGCGCTTGCTAGGG C TAG ATG TA |
| X 7nt | CCCAGTTGAGCGCTTGCTAGGG C TAG ATG T |
| X T2 | CCCAGTTGAGCGCTTGCTAGGG C TT TAG ATG T |
| X T6 | CCCAGTTGAGCGCTTGCTAGGG C TTT TTT TAG ATG T |
| X T12 | CCCAGTTGAGCGCTTGCTAGGG C TTT TTT TTT TTT TAG ATG T |
| XT18 | CCCAGTTGAGCGCTTGCTAGGG C TTT TTT TTT TTT TTT TTT TAG ATG T |
| H | CCGTAGCAGCGCGAGCGGTGGG |
| H 8nt | CCGTAGCAGCGCGAGCGGTGGG C TAG ATG TA |
| H 7nt | CCGTAGCAGCGCGAGCGGTGGG C TAG ATG T |
| H T2 | CCGTAGCAGCGCGAGCGGTGGG C TT TAG ATG T |
| H T6 | CCGTAGCAGCGCGAGCGGTGGG C TTT TTT TAG ATG T |
| H T12 | CCGTAGCAGCGCGAGCGGTGGG C TTT TTT TTT TTT TAG ATG T |

| | |
|---|---|
| H T18 | CCGTAGCAGCGCGAGCGGTGGG C TTT TTT TTT TTT TTT TTT TAG ATG T |
| B | CCCTAGCAAGCCGCTGCTACGG |
| B 8nt | CCCTAGCAAGCCGCTGCTACGG C TAG ATG TA |
| B 7nt | CCCTAGCAAGCCGCTGCTACGG C TAG ATG T |
| B T2 | CCCTAGCAAGCCGCTGCTACGG C TT TAG ATG T |
| B T6 | CCCTAGCAAGCCGCTGCTACGG C TTT TTT TAG ATG T |
| B T12 | CCCTAGCAAGCCGCTGCTACGG C TTT TTT TTT TTT TAG ATG T |
| B T18 | CCCTAGCAAGCCGCTGCTACGG C TTT TTT TTT TTT TTT TTT TAG ATG T |
| 9nt atto655 | Atto655 C TAG ATG TAT |
| 8nt atto655 | Atto655 C TAG ATG TA |
| 8nt atto643 | Atto643 C TAG ATG TA |
| 7nt atto655 | Atto655 C TAG ATG T |
| 12nt atto655 | Atto655 C TAG ATG TAT TAT |
| Receptor strand | SH - CAT CAT CAT ACG CTT CCA AT A ATA CAT CTA |
| FAM-receptor strand | SH - CAT CAT CAT ACG CTT CCA AT A ATA CAT CTA-FAM |
| Non-binding receptor strand | SH-TTT ATA CAT TGA TCC TCC CA |
| Substrate docking (AuNR to substrate docking) | Biotin-TTC TAG ATG TAT TAT |

Gel analysis

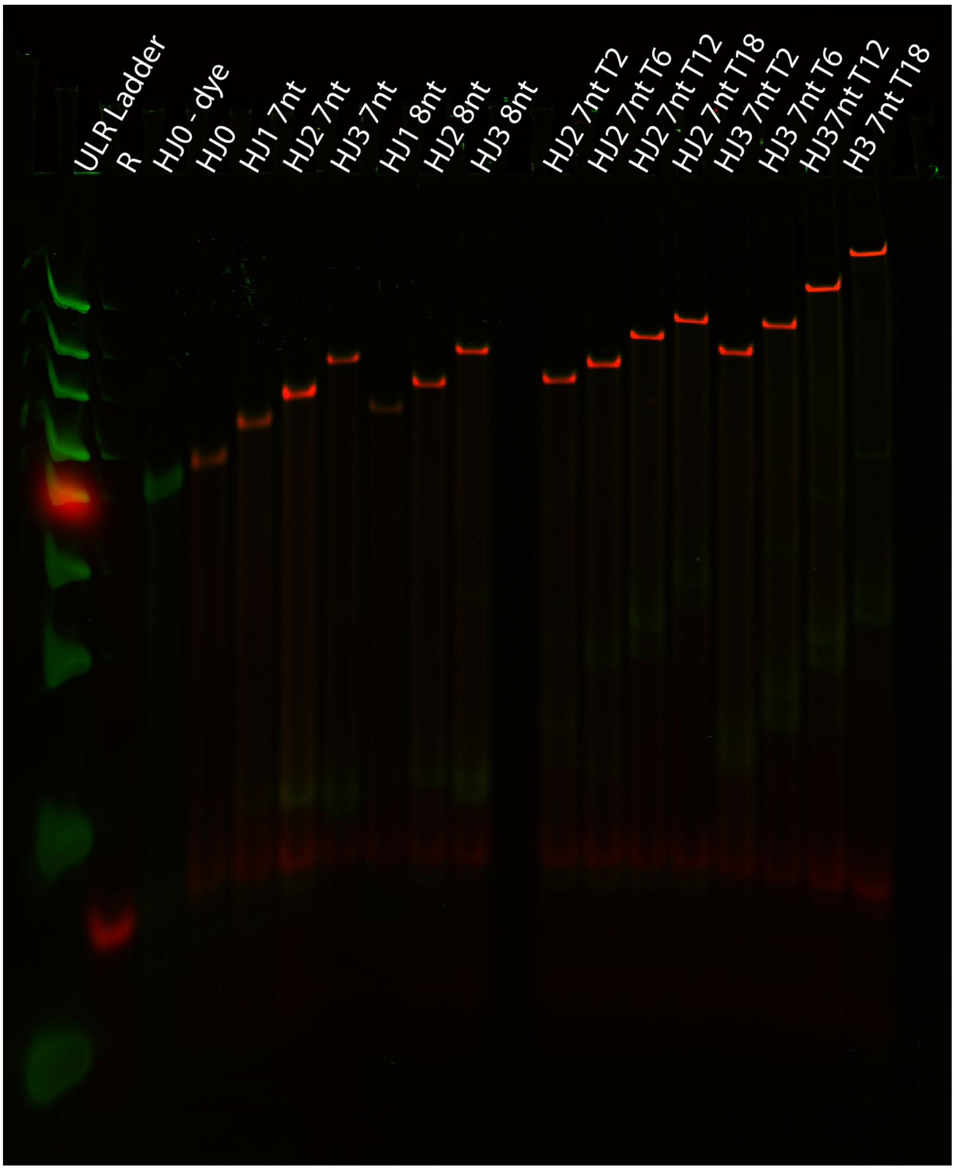

*Figure S2 PAGE gel of HJ constructs. The figure shows a PAGE gel of the assembled HJ contructs. SYBR gold DNA stain is shown in green and ATTO 655 signal in red. Ultralow range DNA ladder is shown on the left with xylene cyanol appearing as a rep spot. The gel shows the expected decreased electrophoretic mobility with added ligand arms and length of ligand sequence.*

#### S2. Determining monovalent on-rates

The ligand sequences are based on a 9nt imager strand that has been used for DNA PAINT on DNA origami[3,4]and gold nanorods[2], with a reported $\mathrm{k_{on}} = 2.3 \cdot 10^6 \mathrm{M^{-1}s^{-1}}$. We use this value as a reference value for the monovalent on-rate $k_{on,M}$ in our model (see section on model derivation). For the 7nt and 8nt ligands we measured the event frequencies relative to the 9nt to get the on-rate of the 7nt and 8nt ligands. We use this way of estimating the on-rates instead of calculating it directly from the event frequency.

The on-rate can be calculated from the event frequency using[5] $F_{event} = N \cdot C \cdot k_{on}$ where N is the number of receptor strands, and C, is the concentration of ligands. Calculating it directly from the event frequency is challenging because it requires that we know the number of receptor strands on each particle. The number of receptor strands may vary because of particle size differences and possibly heterogenous receptor coverage[4]. Our approach was instead to generate time-traces for the 9nt ligand and for the 7nt or 8nt ligands from the same particles by staying on the same field-of-view thereby eliminating the particle-to-particle differences. In this way we compare the event frequencies on each particle for the 9nt to 7nt and 9nt to 8nt ligands. Using a peristaltic pump (which) we could sequentially flow in the two ligands at same concentration while staying on the same field-of-view. Between measuring the first and second ligands we flowed in with buffer until we did not observe any binding events to ensure that no ligands were left in the flow cell. The laser beam was blocked during flow to avoid receptor strand depletion[6]. By comparing the event frequencies, we can estimate the relative on-rate:

$$\frac{\mathrm{k_{on,9nt}}}{\mathrm{k_{on,7/8nt}}} = \frac{\mathrm{F_{9nt}}}{\mathrm{F_{7/8nt}}}$$

As expected[3] we found only small differences for the on-rates: $\frac{\mathrm{k_{on,7nt}}}{\mathrm{k_{on,9nt}}} = 0.95 \pm 0.18$ and $\frac{\mathrm{k_{on,8nt}}}{\mathrm{k_{on,9nt}}} = 1.24 \pm 0.18$.

#### S3. Determining monovalent bound-state lifetimes ($\tau_{on}$)

The monovalent bound-state lifetime is a parameter in the step-binding model. We determine the value from the 7nt and 8nt imagers respectively by measuring the bound-state lifetime values in time traces from three measurements of different FOVs. We extract the average lifetime values by fitting the cumulative distribution function (CDF). For the 7nt monovalent lifetime we found: $\tau_{7nt} = 0.0121\mathrm{s} \pm 4 \cdot 10^{-4}\mathrm{s}$ and for the 8nt ligand $\tau_{8nt} = 0.024\mathrm{s} \pm 0.0012\mathrm{s}$. We have used these values for $k_{on,M}$ in the step binding model ($k_{on} = 1/\tau$).

#### S4. Ligand DNA hairpin formation

We observe a drastic change in bound-state lifetimes when introducing T-spacers in our HJ binding sequences (see fig 5. In main paper) that is not explained by the restriction factor or drop in effective concentration. The drop is most likely due to self-interaction (transient hairpin formation, etc.) in the ligand strand that is known to affect the on-rate in strand hybridization[7].

In order to quantitatively estimate the difference in on-rate between the 7nt and the 7nt with added T-spacers we measured the event frequencies under the same imaging conditions (temperature, concentration, etc.). To avoid the particle-to-particle differences in event frequencies we performed measurements on the same particles for a 7nt and a 7nt+T2 ligans as described in the previous section. We found the measured relative on-rate $\frac{k_{on,7nt+T2}}{k_{on,7nt}} = 0.36 \pm 0.16$. We have used 0.36 to correct the on-rates for all the T-spacer constructs in the model.

In order to investigate whether the ligand strands can potentially self-interact we have used the NUPACK[8] simulation tool. The simulations were performed at 27 deg and with 5mM NaCl (lowest allowed in the software) and 10 mM $MgCl_2$. The 7nt shows some self-interaction, but introducing the T spacers gives rise to a lot stronger interaction between the C and G bases in the sequence. We also see that the equilibrium probability is only weakly changing when adding more than 2 T-spacers. Therefore, we use the T2 correction factor for all the spacer constructs in our model.

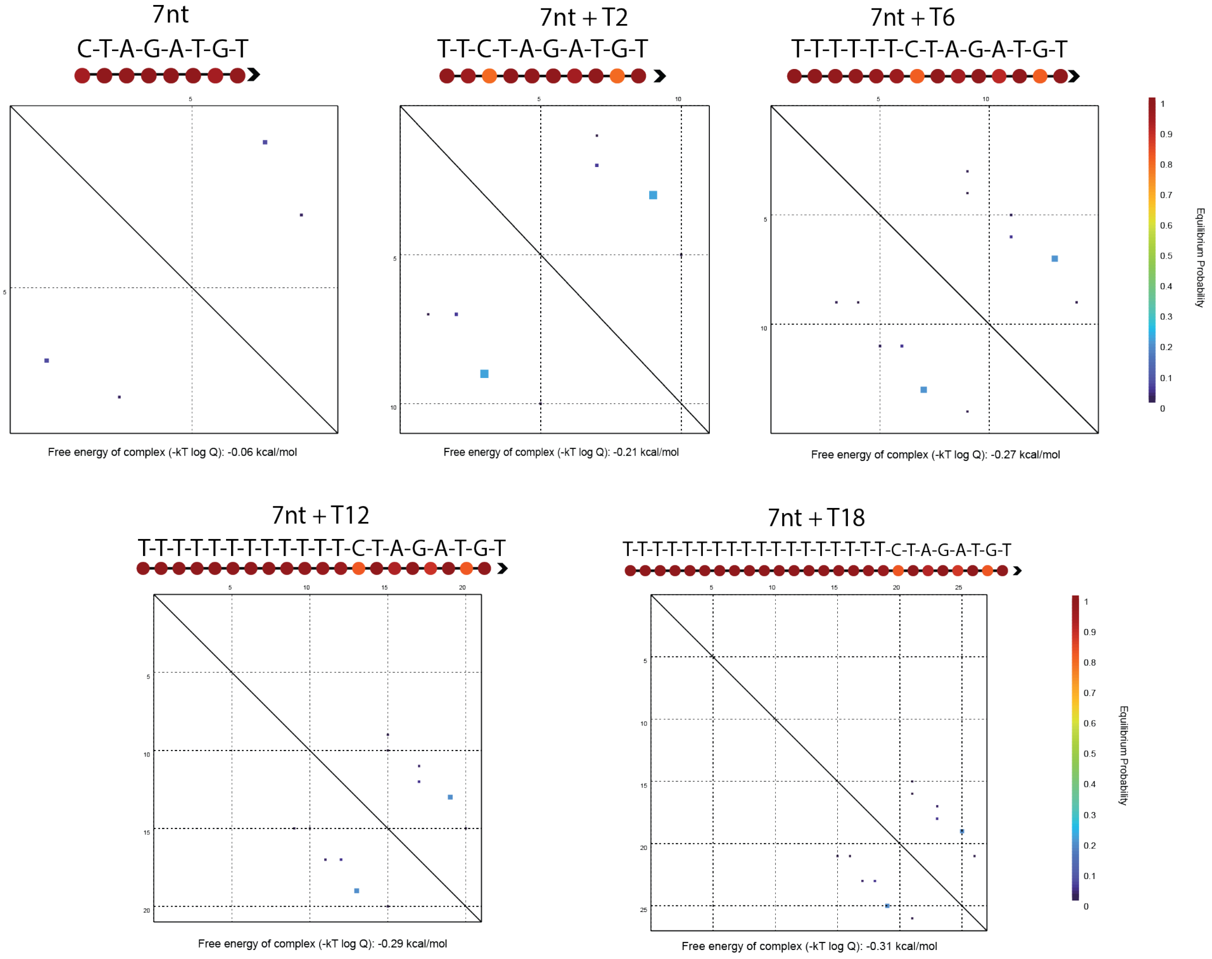

*Figure S3 Nupack analysis of ligand sequences. The figures shows the NUPACK analysis for the 7nt imagers with and without spacers. For each ligand is shown the sequence, the base interaction probability and interaction diagram.*

#### S5. Control for nonspecific binding HJ with no ligand

We have constructed a HJ without binding arms to test for nonspecific binding of the HJ and dye to the particles. In comparison with the monovalent HJ (HJ1) with a 7nt binding sequence we find that $\sim 55\%$ of particles show no events at all compared to 10% for HJ1. Imaging conditions, buffer conditions and concentrations were the same for this comparison. Nonspecific interactions (or HJ diffusing slowly near the particles) do occur but are very infrequent and will therefore only make up a small fraction of our event data for every trace. We observed only $\sim 1.2$ events pr. Particle on average compared to $\sim 4.1$ for HJ1 (those that showed events).

#### S6. Nanorod size measurements and receptor density

##### TEM measurements

We used transmission electron microscopy to measure the length and width of the nanorods

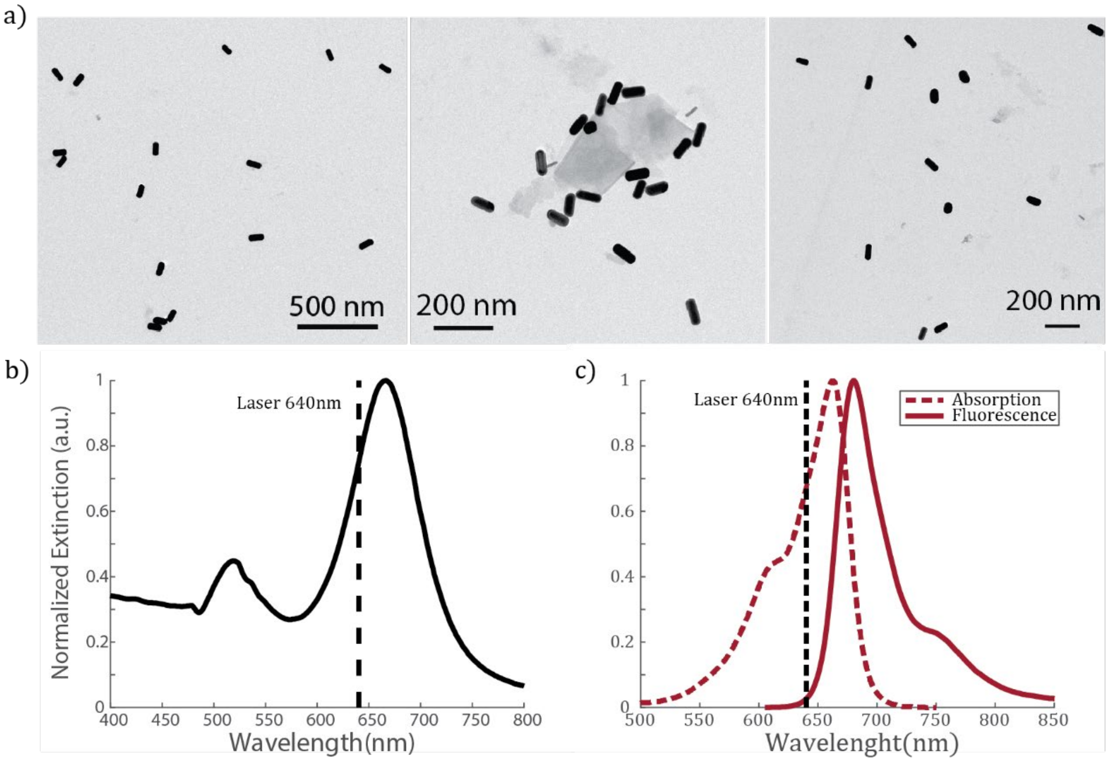

*Figure S4 a) TEM images of Au nanorods. From 50 nanorods the average length and width was determined. L=84.7±9.6nm and W=37.6±4.1nm. b) Normalized extinction spectra of the nanorod suspension used in this work with LSPR around 665, with inset of the 640nm laser shown with the dotted line. C) Absorption and fluorescence spectra (normalized intensity) of the ATTO655 dyes (data from attotec, gmbh www.atto-tec.com)*

##### Measuring DNA loading

In order to estimate the surface density of receptor DNA on the gold nanorods we measured the number of DNA receptor strands attached to gold nanorods in suspension. Here we employed the commonly used procedure of attaching a fluorescently labeled receptor strands and later etching away the nanoparticle to measure the concentration of bound receptor DNA[9]. We used receptor strands with a FAM fluorescent tag on the 3'-end to attach to the gold particles using the freeze-thaw protocol[10]. After freeze-thawing we measured the absorbance and quantified the nanoparticle concentration using a standard curve for absorbance of nanorod suspensions with different known concentrations. Using a commercial gold etchant, we etched the nanorods by adding 1uL to 100uL of our DNA functionalized nanorod suspension. We then measured fluorescence signal from the suspension and derived the DNA concentration from a fluorescence standard curve of different concentrations of the FAM labelled DNA in the same etching buffer as used for the nanorod suspension. From three experimental repeats (three batches of freeze-thaw prepared FAM-DNA functionalized nanorods) we obtained an average value of 1075 DNA strands pr particle with a standard deviation of 352nm. Fluorescence and Absorbance measurements were carried out on a Denovix DS-11 FX+ nanodrop.

#### Estimating receptor density

To estimate the receptor density on the nanorods, we use a similar approach as Horáček, et al[4]. Here, we express the maximum number of DNA on the surface of a rod by taking into account the DNA layer. We consider a layer of closely packed DNA spheres covering the particle and relate the maximum number of bound DNA spheres to the radius of such DNA spheres by taking the ratio of the surface area of the DNA layer $\mathrm{R_{DNA}}$ away from the particle and the footprint of a DNA sphere $A_{DNA}$:

$\mathrm{N_{max}} = \frac{\mathrm{SA}}{\mathrm{A_{DNA}}}$

Since we approximate the DNA strands as spheres there is additional void area to be considered. We do this by adding the packing density of equal circles $\eta \approx 0.91$.

$$\mathrm{N_{max}} = \frac{\mathrm{SA}}{\mathrm{A_{DNA}}} \cdot \eta$$

The nanorod geometry can be approximated as a cylinder with spherically capped ends. In Figure S5 is shown all relevant parameters.

$$\mathrm{N_{max}} = \frac{2\left(\mathrm{R_{DNA}} + \mathrm{R_p}\right)\mathrm{D} + 4(\mathrm{R_{DNA}} + \mathrm{R_P})^2}{\mathrm{R_{DNA}^2}} \cdot \eta$$

$R_p$ is the radius of the spherical ends of the rod, i.e. half of the width, $\mathrm{R_p} = \frac{\mathrm{W}}{2}$, D is the length of the cylindrical part, i.e. the length of the rod minus the radius of the spherical ends $\mathrm{D} = \mathrm{L} - 2\mathrm{R_P}$, and $R_{DNA}$is the radius of the DNA spheres. Using the measured average particle length and widths and the measured average DNA loading as $\mathrm{N_{max}}$ we get $\mathrm{R_{DNA}} \approx 1.75\mathrm{nm}$.
Still assuming a closed packing of the circles, the lattice parameter a is twice the radius of a DNA sphere: $2\mathrm{R_{DNA}} = \mathrm{a} \approx 3.50\mathrm{nm}$
The surface coverage, $\Gamma_s$ (in moles pr. Area), in a closed packed system can be expressed by the lattice parameter: $\Gamma_s = \frac{2}{\sqrt{3}\, a^2 N_{av}}$ . Using the estimated values of the lattice parameter we find a surface coverage of $1.6\ \mathrm{mol\ cm^{-2}}$.

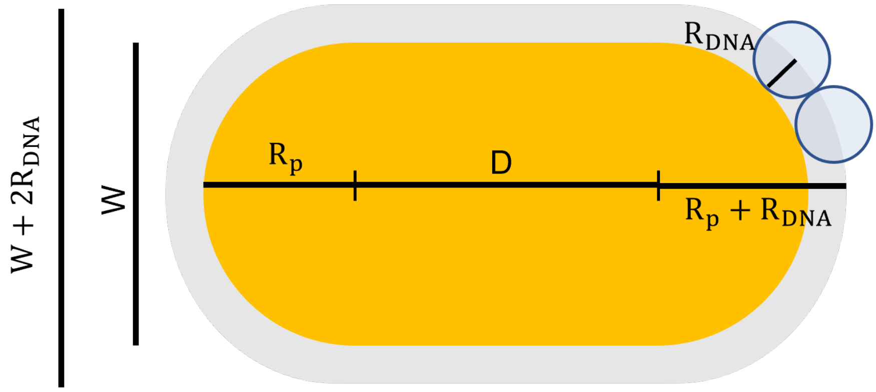

*Figure S5* Illustration of nanorod with relevant parameters.

#### Immobilizing nanorods on functionalized glass surface

The gold nanorods are attached to the surface through a BSA-biotin that is bound to the surface of the glass (Figure S6). Neutravidin bridges the BSA-biotin and biotinylated substrate-docking DNA that is complimentary to the receptor strands on the particles.  In order to check the ratio of single particles to dimer or aggregates we did scanning electron microscopy (Magellan SEM, FEI) on a used sample. We dried the sample by first flushing it with 96% ethanol (sigma) and then air-dried. Then we released part of the glass in the ibidi channels (our functional sample site) and evaporated a 5nm layer of Ti using electron assisted physical vapor deposition (e-pvd, cryofox, polyteknik) for better SEM imaging. We found that $\sim 80\%$ were single particles dispersed on the functionalized glass surface. In Figure S6 we show two exemplary SEM images with single particles highlighted with green circles and dimer/aggregates with red.

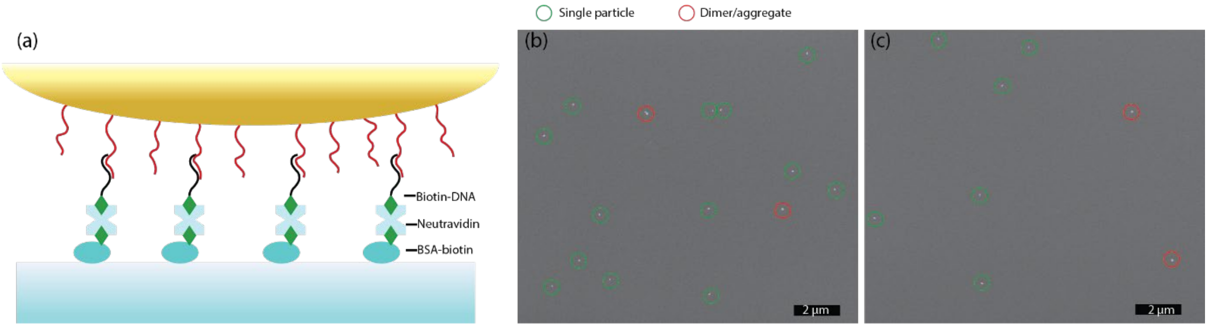

*Figure S6 AuNR on glass surface. a) Sketch of the surface functionalization for AuNR immobilization on the glass surface. b) and c) show SEM images of glass surface immobilized nanorods. Green circles represent single nanorods and red circles represent dimers or aggregates.*

### S7. Modelling bound-state lifetimes ($\tau_{on}$)

#### Step-binding model

Following the work of Huskens, et. al. we use a step-binding model[11] to estimate the off-rate of the multivalent binding HJ (see figure 3 and 4 in the main paper). The first step is the binding/unbinding of the HJ to the surface which is determined by the off-rate and the on-rate of the single ligand ($k_{on,M}$ and $k_{off,M}$ ) with an additional statistical factor because the HJ carries 2 or 3 binding arms. The second step is the binding/unbinding of the second arm while bound with the first arm. The rates are again determined by the single ligand rates but the on-rate is furthermore determined by the effective concentration $C_{eff}$. For the 3 arm HJ the third step is the binding/unbinding of the third arm while bound with two. The third on-rate is now modified by the effective concentration $C_{eff}$ of this third step which is different from the second step. Using the step model we can express the multivalent on-rate in terms of the effective concentrations and the monovalent rate constants.

Starting with the bivalent case (two armed) we can use the above scheme and write a bivalent equilibrium constant (association constant) $K_{eq,B}$:

$$K_{eq,B} = \frac{2k_{on,M} \cdot k_{on,M} \cdot C_{eff}}{k_{off,M} \cdot 2k_{off,M}}$$

We assume that the on-rate of the bivalent is 2 times the monovalent on-rate:

$$k_{on,B} = 2k_{on,M}$$

And we can write the bivalent equilibrium in terms of the divalent on and off-rates:

$$K_{eq,B} = \frac{k_{on,B}}{k_{off,B}}$$

Combining the three equations we obtain the following expression:

$$k_{off,B} = \frac{2\ k_{off,M}}{K_{eq,M} \cdot C_{eff}}$$

Which gives the bound-state lifetime:

$$\tau_{on,B} = \frac{K_{eq,M}\ C_{eff}}{2\ k_{off,M}}$$

We can express the bound-state lifetime for the HJ with 3 binding arms with the same approach as for the 2-armed HJ. In this case there is an additional binding step when the third arm binds with an additional effective concentration, $C_{eff,B\rightarrow T}$ associated with the third arm binding. We can then write up the three equations describing the equilibrium constant and binding constant for the 3 armed HJ:

$$K_{eq,T} = \frac{3k_{on,M}\ 2k_{on,M}\ C_{eff,M\rightarrow B}\ k_{on,M}\ C_{eff,B\rightarrow T}}{k_{off,M}\ 2k_{off,M}\ 3k_{off,M}}$$

$$k_{on,T} = 3k_{on,M}$$

$$K_{eq,T} = \frac{k_{on,T}}{k_{off,T}}$$

Combining the three equations we obtain the following expressions for the rate and bound-state lifetime respectively:

$$k_{off,T} = \frac{3\ k_{off,M}}{K_{eq,M}^2\ C_{eff,M\rightarrow B}\ C_{eff,B\rightarrow T}}$$

$$\tau_{on,T} = \frac{K_{eq,M}^2\ C_{eff,M\rightarrow B}\ \ C_{eff,B\rightarrow T}}{3\ k_{off,M}}$$

#### Effective concentration

The effective concentration is an important parameter when describing multivalent systems binding at surfaces (and in solution as well) [12]. When bound to the surface the unbound ligands on the multivalent object are confined to a small volume close to the receptor surface. This gives rise to a high local concentration of ligands and receptors that increase the binding probability of the subsequent ligands after the first ligand-receptor binding. In the following we try to estimate the effective concentration by taking into account the HJ and the DNA surface on the gold particles. The effective concentration $C_{eff}$ is related to the unrestricted probing volume of the second binding arm. It is the number of available host sites, $n_h$, in an area underneath the probing volume, $(l)$, divided by the probing volume itself[12]:

$$C_{eff} = \frac{n_h(L)}{N_{av}\ V(L)}$$

The number of available host sites is given by the surface area, A, the molar surface coverage, $\Gamma_s$, and the Avogadro constant $N_{av}$:

$$n_h(L) = A\ N_{av}\ \Gamma_s$$

The probing volume is the volume which the ligand can occupy while bound to the surface through a receptor-ligand bond (DNA-DNA bond). We can describe the volume in multiple ways taking into account the geometry and structure of the HJ construct and the receptor surface geometry. In the work of Huskens et al[11], that we build upon, the volume is considered to be a free diffusing ligand constrained to a volume defined by the polymer that tethers it to the surface. In the bivalent case on a *flat receptor surface*, the volume is a hemisphere with radius equal to the end-to-end distance between two ligands.

$$V = \frac{2}{3}\pi L_{ete}^3$$

And the area on the surface is then:

$$A = \pi L_{ete}^2$$

The HJ is built up of 4 arms where each arm can carry a single stranded DNA (ssDNA) part that includes a spacer sequence with a C nucleotide (and additional multiple T nucleotides) and the ligand sequence which is 8 or 7 nucleotides. We divide the construct into different segments when bound with the first ligand (Figure S7): $\mathrm{L_{spacer}}, \mathrm{L_{HJ}}, \text{and } \mathrm{L_{spacer+ligand}}$.
In the following we estimate the lengths of the single stranded DNA parts using a worm-like-chain model approach to compute the end-to-end lengths[13]:

$$\mathrm{L_{ete}} = \sqrt{2 \cdot \mathrm{P} \cdot (\mathrm{l_s} - \mathrm{P} + \mathrm{P} \cdot \exp(-\mathrm{l_s}/\mathrm{P}))}$$

Where P is the persistence length of single stranded DNA and $l_s$ is the segment length i.e., the length of a single nucleotide. Both the persistent length and segment length is dependent on salt concentration, we use P=2nm and $\mathrm{l_s} = 0.63\mathrm{nm}$ in our calculations [14].
We set the end to end length of the HJ in the bivalent case as $L_{HJ} = 7.5\ nm$ since the two ligands are spanned by 22 nucleotides and the distances between nucleotides in double stranded form is $\sim 0.34nm$.

In the case of the HJ used in our study we can include the structure of the HJ as well.
In the *unstructured* case we consider it to be a free diffusing ligand whereas for the *structured* we attempt to take into account the structure of the HJ (see figure3 main paper).
The HJ arms can stack in two distinct isoforms that are similar to an X-like structure under the used buffer conditions[2]. For the bivalent construct, the two ligand strands are placed such that they are always at opposite ends (far apart) of the HJ, which means they are always separated by the span of the double stranded HJ. We approximate the HJ in the bivalent case as a rigid segment between the two ligands. Furthermore, the HJ is connected to the surface through a ligand and we consider the length of the HJ to start from above the double stranded binding to the receptor strand, i.e. from the spacer part of the ligand sequence. The HJ can then swing around from the surface in a hemisphere. The ligand sequence on the unbound arm can rotate around freely. It can span a sphere around the point to which it is connected to the HJ. In that way the probing volume forms a hemi-spherical shell. The outer radius is the length of the spacer on the bound arm, the span of the HJ, the length of the spacer on the unbound arm, and the length of the ligand region, as it has to reach with its outermost nucleotide for binding. The inner radius is the length of the spacer on the bound arm and the span of the HJ minus the length of the spacer and ligand on the unbound arm.

$$V_{structured} = \frac{2}{3} \cdot \pi \cdot (r_2^3 - r_1^3)$$
$$= \frac{2}{3} \cdot \pi \cdot \left( \left(L_{HJ} + L_{spacer} + L_{ligand+spacer}\right)^3 - \left(L_{spacer} + L_{HJ} - L_{spacer+ligand}\right)^3 \right)$$

The area under the probing volume is then spanned by two circles of the same radii as for the shell. Here we neglect the fact that the ssDNA part of the bound arm is not rigid.

$$A_{structured} = \pi \cdot (r_2^2 - r_1^2)$$
$$= \pi \cdot \left( \left( L_{HJ} + L_{spacer} + L_{ligand+spacer} \right)^2 - \left( L_{spacer} + L_{HJ} - L_{spacer+ligand} \right)^2 \right)$$

In our *unstructured* model the hemispherical volume is determined by L which is the same outer L in the *structured* model, which gives the following volume and area:

$$V_{unstructured} = \frac{2}{3}\pi \left( L_{HJ} + L_{spacer} + L_{ligand+spacer} \right)^3$$
$$A_{unstructured} = \pi \left( L_{HJ} + L_{spacer} + L_{ligand+spacer} \right)^2$$

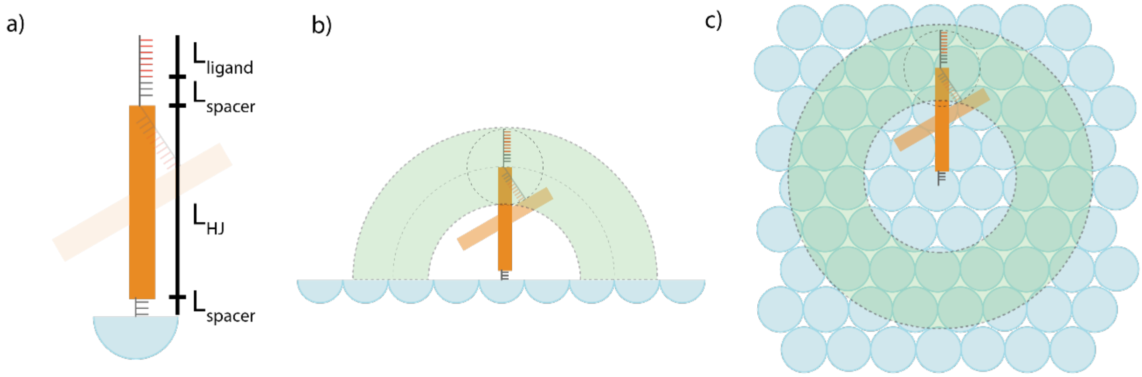

*Figure S7 Sketch of the HJ parameters in estimating Ceff. a) HJ represented as a rod, when binding with the ligand far apart, with the different length parameters used in the Ceff estimation. The HJ is bound to a receptor DNA strand (blue halfsphere). A spacer sequence (the C nucleotide and the possible T spacers) is separating the bound ligand and the rigid HJ part with lengths $L_{spacer}$ and $L_{HJ}$ respectively. The second ligand arm is spanned by the spacer and the binder, $L_{ligand}$. b) and c) shows the sideview (b) and topview (c) of the hemispherical shell for the structured model on a flat surface.*

In our case the surface is a nanorod and not a flat surface. Therefore, we must consider the effect of the curved surface on the probing volume and area. In this case the volume is now larger than for a flat surface. We consider the situation of the spherical part (end parts of the rod) to account for the curvature. The probing volume can be split into two parts as shown in Figure S8. The red part is the hemispherical volume that is the same as for the flat surface. The additional volume (blue) that comes from the curvature of the particle can be considered as the segment of a sphere (Figure S8B) minus the volume of the spherical cap on the particle sphere shown in green.

The radius of the top sphere in Figure S8 comes from the arc length that we define as the length of the construct $\left(L_{HJ} + L_{spacer} + L_{binder+spacer}\right)$. This gives a slightly smaller radius for the blue sphere. The radius is the length of the line C that can be extracted from the isosceles triangle in Figure S8B; $\mathrm{C} = 2 \cdot \mathrm{R_p} \cdot \cos\alpha$. Where $\alpha = \frac{180° - \theta}{2}$, $\theta = \frac{L}{R_p} \cdot \frac{180°}{\pi}$, and $L = L_{HJ} + L_{spacer} + L_{binder+spacer}$.

The volume of the segment is defined by the radii of the two segmentation lines $r_1$ and $r_2$, where $r_1$ is the radius of the top sphere equal to C, and $r_2$ is the radius of the bottom segmentation line that is defined from the crossing point of the blue sphere and the particle sphere. From the right triangle we can get $r_2$ and the height of the spherical cap h:

$r_2 = \sqrt{C^2 - h^2}$ and $h = C \cdot \cos\alpha$.

The volume of the spherical segment and the cap are:

$$V_{segment} = \frac{\pi h}{6}(3r_1^3 + 3r_2^3 + h^2) \text{ and } V_{cap} = \frac{\pi h}{6}(3r_2^3 + h^2)$$

The volume highlighted in blue is then:

$$V_2 = V_{segment} - V_{cap}$$

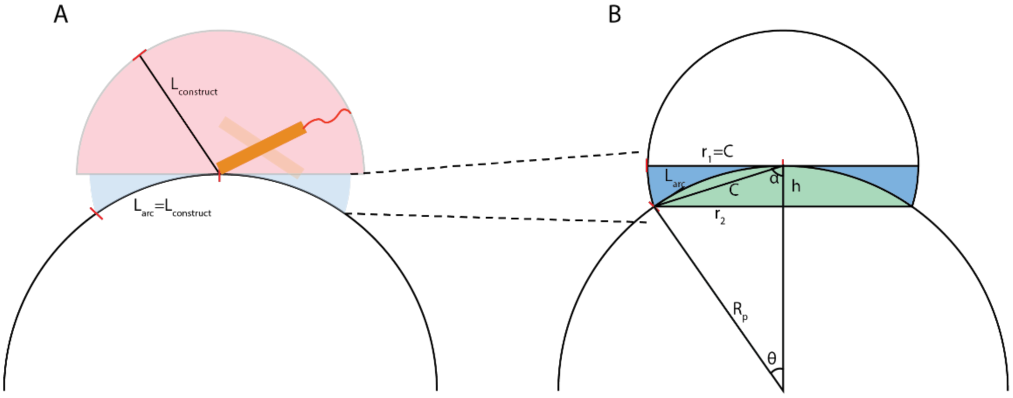

*Figure S8 Schemtics of the Probing volume. a) The probing volume can be divided into two sections. A half sphere (red) where the radius is the length of the HJ construct and a spherical section (blue) with overlapping particle sphere (white) determined by the arc length which is the length of the HJ construct. (b) showing the dimensions of the spherical segment and the particle overlap in green. Here is shown the isosceles triangle used to calculate the volume.*

The area that the HJ can probe is the area of the spherical cap in green, which is given by

$$A = \pi(r_2^2 + h^2)$$

In our case the radius of the particle sphere is the measured particle radius (W/2) and the DNA layer. We consider the DNA layer thickness from the thiol bond on the particle surface to the first nucleotide in the complementary sequence which is then 22-23 nucleotides for 8 and 7nt ligand sequences respectively. We have used the length of 22 nucleotides in our calculations.
For the *structured* model, we again define an inner and outer radius for the spherical shell using $L_{outer} = L_{HJ} + L_{spacer} + L_{ligand+spacer}$ and $L_{inner} = L_{spacer} + L_{HJ} - L_{spacer+ligand}$.

In order to estimate the effective concentration in the third binding step for a 3-armed HJ, we use the same HJ geometry for the second step in the trivalent binding as for the bivalent HJ, i.e. we say that in the second binding step it binds with the far apart ligand. Then in the third step we assume that during the second step the HJ is in close contact with the surface. With that assumption we describe the probing volume for the third ligand as a sphere spanned by the length of the spacer and the ligand. Then we use the following expressions for the Area and volume in the third binding step.

$$V = \frac{2}{3} \cdot \pi \cdot r^3 = \frac{2}{3} \cdot \pi \cdot \left(L_{spacer} + L_{binder}\right)^3$$
$$A = \pi \cdot r^2 = \pi \cdot \left(L_{spacer} + L_{binder}\right)^2$$

With addition of T-spacers we can estimate the effect on $C_{eff}$. In Figure S9is shown the $C_{eff}$ with added T-spacer and we see a clear drop for both the models in the bivalent case and for the structured model in the trivalent case, which is directly related to the increase in $L_{spacer}$.

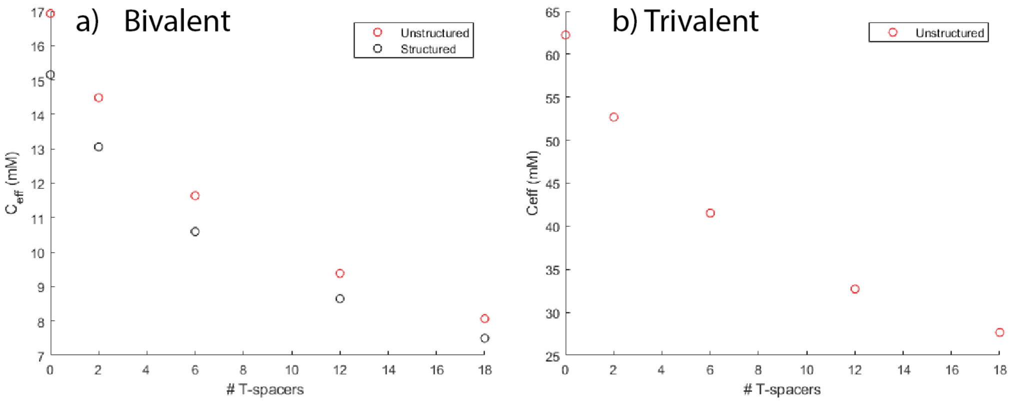

*Figure S9 Calculated effective concentrations ($C_{eff}$) of a bivalent (a) and trivalent (b) HJ with a 7nt ligand with added T-spacers. In red is shown the unstructured model values and in black the structured model values as described in the text. The $C_{eff}$ drops with the addition of T spacer, i.e. increasing the length of the ligands.*

#### S8. Transition rates

The step binding model allows us to estimate the average transition rate within each step. We have listed the transition rates going from monovalent to bivalent and from bivalent to monovalent in Table S2, and the transition rates going from bivalent to trivalent and from trivalent to bivalent in Table S3. We see that the monovalent to bivalent is by far the fastest transition.

*Table S2 list of transition rates from monovalent to bivalent and bivalent to monovalent.*

| $M \leftrightarrow B$ | $2k_{on,M} \cdot C_{M\to B}^{free} \cdot \omega_{M\to B}$ | $2k_{off}$ |
|---|---|---|
| 7nt | $1.04 \cdot 10^4 s^{-1}$ | $167\ s^{-1}$ |
| 8nt | $1.43 \cdot 10^4 s^{-1}$ | $83 s^{-1}$ |

*Table S3 list of transition rates from bivalent to trivalent and trivalent to bivalent.*

| $B \leftrightarrow T$ | $k_{on,M} \cdot C_{B\to T}^{free} \cdot \omega_{B\to T}$ | $3k_{off}$ |
|---|---|---|
| 7nt | $813 s^{-1}$ | $250\ s^{-1}$ |
| 8nt | $707 s^{-1}$ | $125\ s^{-1}$ |

### S9. Fluorescent enhancement factors from burst analysis

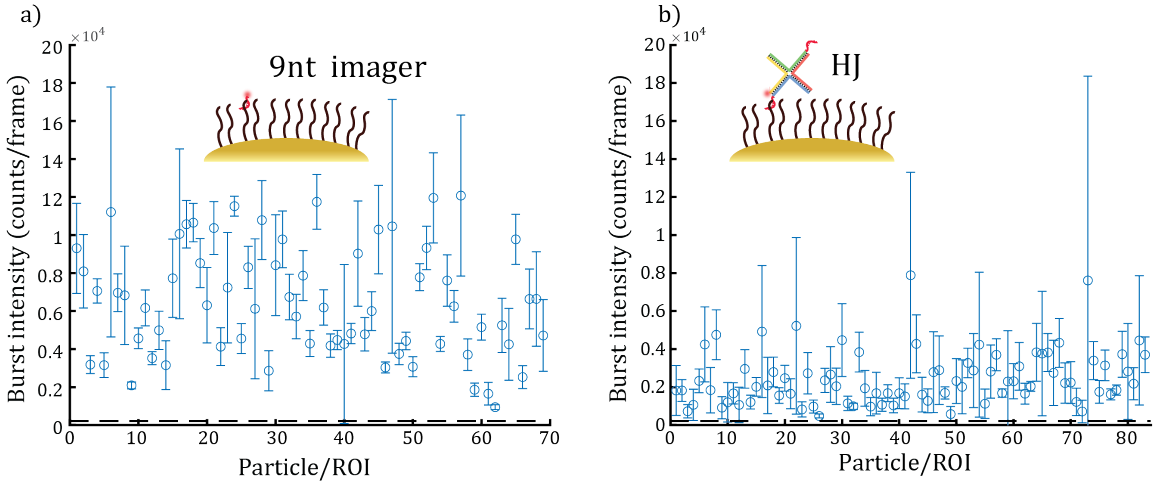

*Figure S10 Fluorescence burst intensity of the 10 most intense bursts from each particle representing binding at the tips. a) From a 9nt imager and b) from a 2-armed HJ, both with 50 ms frames and 20mW laser power. The dotted line represents the average fluorescence intensity from the dye binding in the background, i.e. the nonenhanced fluorescence. For a) the nonenhanced value was* $2.1 \cdot 10^3 \left(\frac{counts}{frame}\right)$ *and for b) it was* $1.5 \cdot 10^3 \left(\frac{counts}{frame}\right)$ *yielding enhancement factors up to* $\sim 58$ *and* $\sim 54$ *for a) and b) respectively.*

The fluorescent enhancement factor is estimated by the fluorescent burst intensity of the 10 most intense bursts for each particle in a timetrace which are shown in Figure S10. The nonenhanced value is found by averaging the burst intensities from dyes binding nonspecifically to the background. This was done by picking 100 random ROIs in the background (not on the particle ROIs) and detect events above threshold. All burst intensities from the background were averaged to the values in the figure caption.

## S10. HJ2 XB

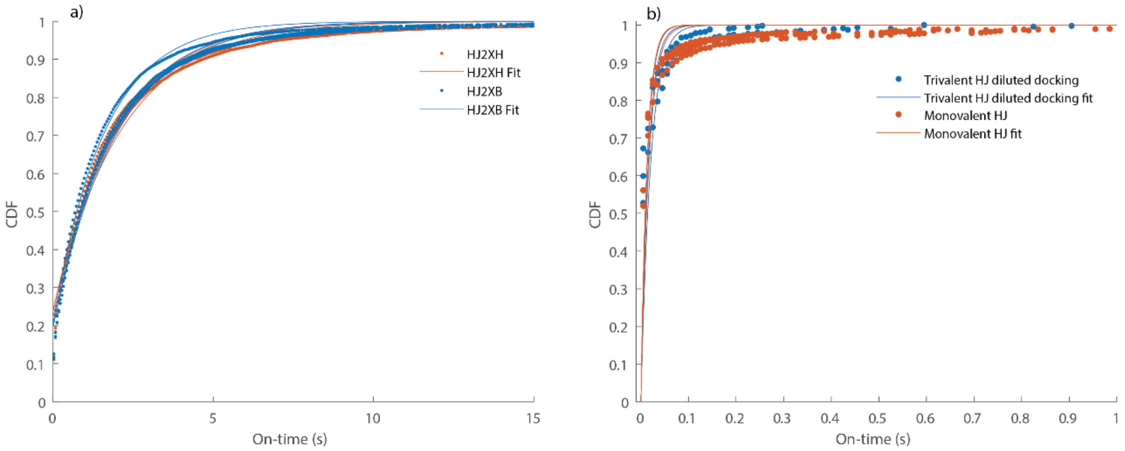

*Figure S11 CDF of bound-state lifetimes (on-times). a) Comparison of 2 bivalent constructs with ligand sequences on XH arms (orange) and XB arms (blue). For each HJ is plotted three CDFS from all time traces within 3 different FOV. The CDFs look almost identical for all 6 CDFs. b) comparison of CDFS from a trivalent HJ binding to particles covered in a 1:50 ratio of receptor DNA and non-binding DNA (blue) and a monovalent HJ binding to particles fully covered in receptor DNA (orange).*

We have constructed two bivalent HJs with ligands placed on different arms. One construct, that has been used throughout the main paper, with ligands placed on XH arms and one with ligands on XB arms. When switching between the two isoform states the binding arms will always bar far apart for the XH construct, whereas it can be close and far apart for the two states respectively for the XB construct. We compare the bound-state lifetimes of the two constructs by plotting the cumulative distribution of binding events from the two bivalent holliday junctions(XH in orange and XB in blue) in Figure S11. From the CDFs it is clear that they show similar behavior and kinetics. This indicates that the far apart binding is preferred, since the close binding would result in a much higher effective concentration that should give rise to longer binding which is not observed.

#### S11 Trivalent HJ binding to sparsely coated particles reduces avidity

As a control to make sure that the increased bound-state lifetime for the trivalent (and bivalent) HJ is indeed due to multivalent binding, we made measurements on trivalent HJs binding to AuNRs covered in sparse density of receptor DNA, such that the HJ can only bind with one ligand at a time. We prepared the AuNRs by freeze-thawing with a 1:50 ratio of non-complementary (non-binding) DNA to receptor DNA, thereby making a surface that is sparsely covered with receptors. In Figure S11 we have plotted the CDF from three measurements from separate FOVs of the trivalent HJ (blue) and compared to a monovalent HJ binding to particles fully covered with receptor DNA (orange). We see that the CDFs are nearly identical meaning that with a sparsely covered surface the trivalent HJ can only bind with one interaction. This shows that the enhanced bound-state lifetimes observed for the multivalent HJs on densely coated particles is a multivalency effect.

#### S12. List of bound-state lifetimes

*Table S4 List of experimental bound-state lifetime s. Each lifetime value is extracted as the average of three measurements from three FOVs. * marks those that were fitted with A=1 and with first data point excluded from the fit.*

| **Sample** | **Bound-state lifetimes (s)** |
|---|---|
| 7nt* | $0.0121 \pm 4 \cdot 10^{-4}$ |
| 8nt* | $0.024 \pm 0.001$ |
| HJ3 8nt | $11.6 \pm 1.90$ |
| HJ2 8nt | $2.06 \pm 0.109$ |
| HJ1 8nt* | $0.0414 \pm 0.003$ |
| HJ2 8nt XB | $1.83 \pm 0.20$ |
| HJ3 7nt no spacer | $2.45 \pm 0.114$ |
| HJ3 7nt T2 | $0.885 \pm 0.059$ |
| HJ3 7nt T6 | $0.875 \pm 0.112$ |
| HJ3 7nt T12 | $0.558 \pm 0.049$ |
| HJ3 7nt T18 | $0.459 \pm 0.036$ |
| HJ1 7nt * | $0.014 \pm 0.002$ |
| HJ2 7nt no spacer | $0.378 \pm 0.043$ |
| HJ2 7nt T2 | $0.240 \pm 0.054$ |
| HJ2 7nt ST2 | $0.097 \pm 0.009$ |
| HJ2 7nt ST6 | $0.077 \pm 0.004$ |
| HJ2 7nt T12 | $0.048 \pm 0.004$ |
| HJ2 7nt T18 | $0.041 \pm 0.0103$ |
| HJ3 8nt 1:50 diluted* | $0.016 \pm 0.004$ |

### S13. Examples of events with 12nt imager using 100 fps framerate

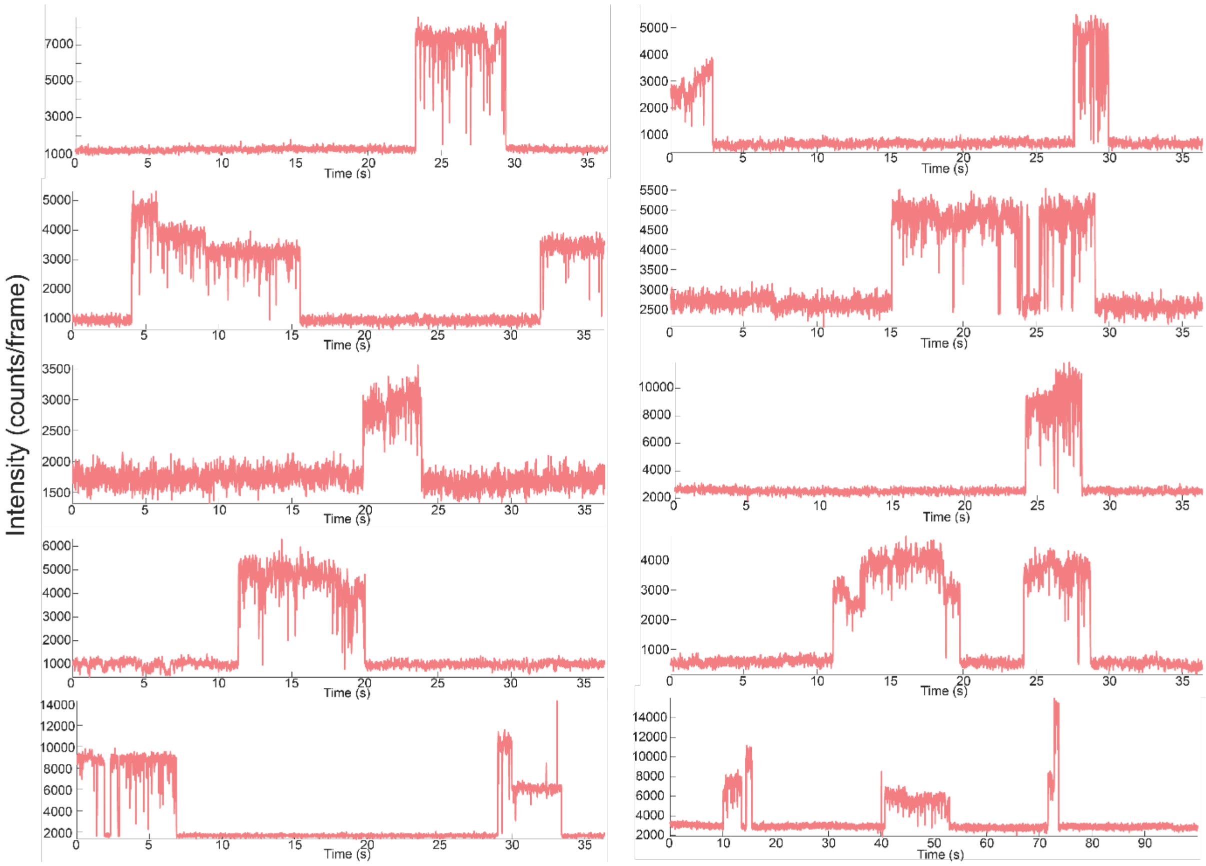

*Figure S12 Examples of binding events form the monovalent 12nt ligand*

### S14. Examples of events with HJ3 8nt using 100 fps framerate

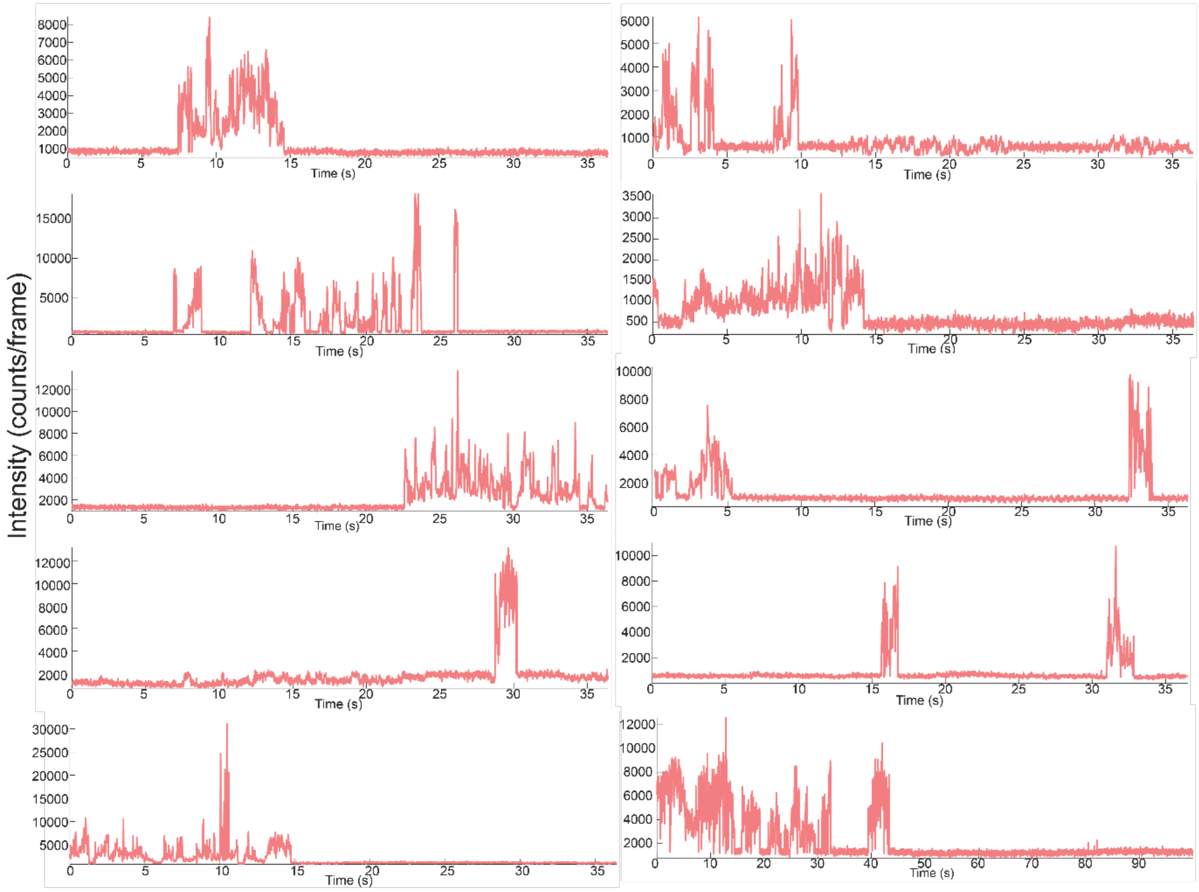

*Figure S13 Examples of trivalent HJ binding events with 100 fps*

#### S16. Examples of events with HJ3 8nt using 1000 fps framerate

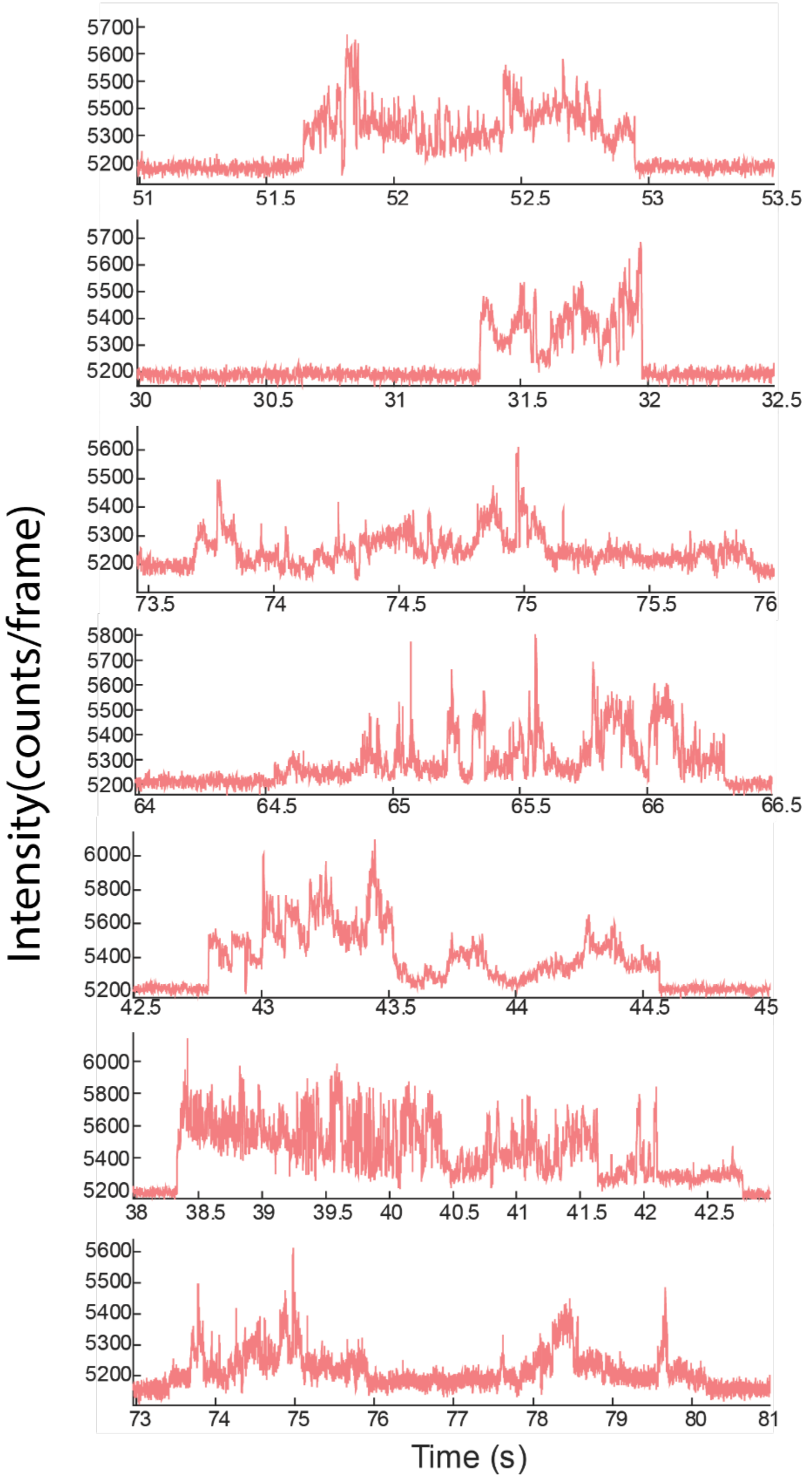

*Figure S14 Examples of Trivalent HJ binding events with 1000 fps acquisition*

#### S17. Simulation points

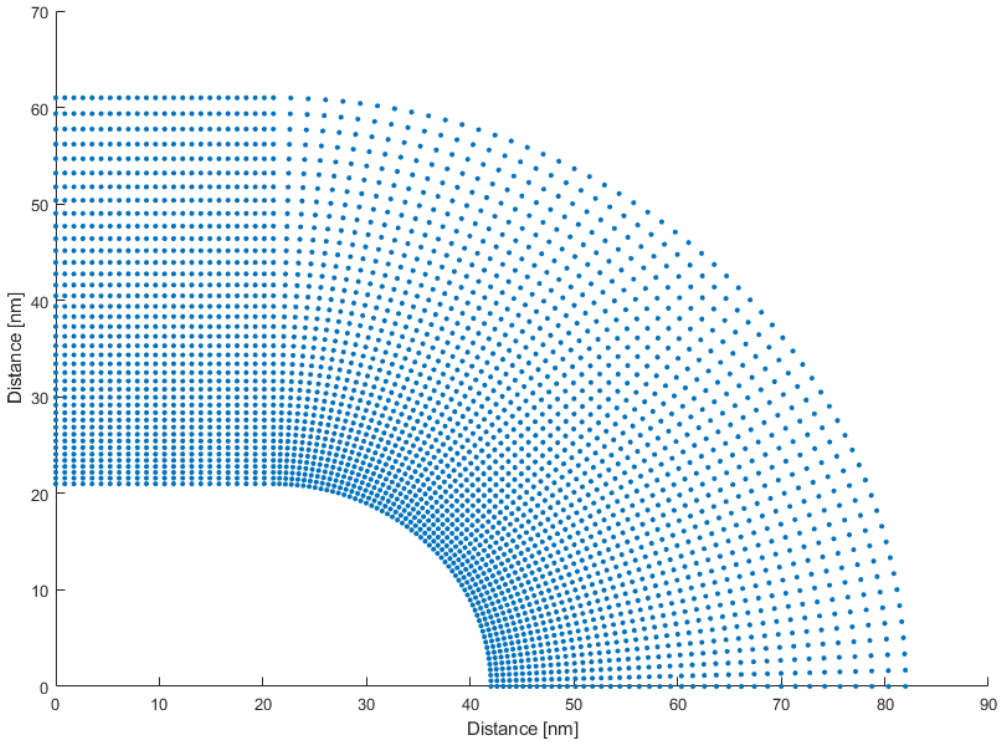

*Figure S15 Simulation points. The figure shows all the points around the nanoparticle where we simulated the fluorescence enhancement using the Boundary Element method and that were later 2D extrapolated to generate the fluorescence enhancement map. For symmetry reasons, it is sufficient to simulate one quarter of the 2D rod.*

Bibliography:

(1) Grainger, R. J.; Murchie, A. I. H.; Lilley, D. M. J. Exchange between Stacking Conformers in a Four-Way DNA Junction. *Biochemistry* **1998**, *37* (1), 23–32. https://doi.org/10.1021/bi9721492.

(2) McKinney, S. A.; Déclais, A.-C.; Lilley, D. M. J.; Ha, T. Structural Dynamics of Individual Holliday Junctions. *Nat. Struct. Biol.* **2003**, *10* (2), 93–97. https://doi.org/10.1038/nsb883.

(3) Jungmann, R.; Steinhauer, C.; Scheible, M.; Kuzyk, A.; Tinnefeld, P.; Simmel, F. C. Single-Molecule Kinetics and Super-Resolution Microscopy by Fluorescence Imaging of Transient Binding on DNA Origami. *Nano Lett.* **2010**, *10* (11), 4756–4761. https://doi.org/10.1021/nl103427w.

(4) Horáček, M.; J. Engels, D.; Zijlstra, P. Dynamic Single-Molecule Counting for the Quantification and Optimization of Nanoparticle Functionalization Protocols. *Nanoscale* **2020**, *12* (6), 4128–4136. https://doi.org/10.1039/C9NR10218C.

(5) Jungmann, R.; Avendaño, M. S.; Dai, M.; Woehrstein, J. B.; Agasti, S. S.; Feiger, Z.; Rodal, A.; Yin, P. Quantitative Super-Resolution Imaging with qPAINT. *Nat. Methods* **2016**, *13* (5), 439–442. https://doi.org/10.1038/nmeth.3804.
(6) Blumhardt, P.; Stein, J.; Mücksch, J.; Stehr, F.; Bauer, J.; Jungmann, R.; Schwille, P. Photo-Induced Depletion of Binding Sites in DNA-PAINT Microscopy. *Molecules* **2018**, *23* (12), 3165. https://doi.org/10.3390/molecules23123165.
(7) Schueder, F.; Stein, J.; Stehr, F.; Auer, A.; Sperl, B.; Strauss, M. T.; Schwille, P.; Jungmann, R. An Order of Magnitude Faster DNA-PAINT Imaging by Optimized Sequence Design and Buffer Conditions. *Nat. Methods* **2019**, *16* (11), 1101–1104. https://doi.org/10.1038/s41592-019-0584-7.
(8) Zadeh, J. N.; Steenberg, C. D.; Bois, J. S.; Wolfe, B. R.; Pierce, M. B.; Khan, A. R.; Dirks, R. M.; Pierce, N. A. NUPACK: Analysis and Design of Nucleic Acid Systems. *J. Comput. Chem.* **2011**, *32* (1), 170–173. https://doi.org/10.1002/jcc.21596.
(9) Demers, L. M.; Mirkin, C. A.; Mucic, R. C.; Reynolds, R. A.; Letsinger, R. L.; Elghanian, R.; Viswanadham, G. A Fluorescence-Based Method for Determining the Surface Coverage and Hybridization Efficiency of Thiol-Capped Oligonucleotides Bound to Gold Thin Films and Nanoparticles. *Anal. Chem.* **2000**, *72* (22), 5535–5541. https://doi.org/10.1021/ac0006627.
(10) Liu, B.; Liu, J. Freezing Directed Construction of Bio/Nano Interfaces: Reagentless Conjugation, Denser Spherical Nucleic Acids, and Better Nanoflares. *J. Am. Chem. Soc.* **2017**, *139* (28), 9471–9474. https://doi.org/10.1021/jacs.7b04885.
(11) Gomez-Casado, A.; Dam, H. H.; Yilmaz, M. D.; Florea, D.; Jonkheijm, P.; Huskens, J. Probing Multivalent Interactions in a Synthetic Host–Guest Complex by Dynamic Force Spectroscopy. *J. Am. Chem. Soc.* **2011**, *133* (28), 10849–10857. https://doi.org/10.1021/ja2016125.
(12) Huskens, J.; Mulder, A.; Auletta, T.; Nijhuis, C. A.; Ludden, M. J. W.; Reinhoudt, D. N. A Model for Describing the Thermodynamics of Multivalent Host−Guest Interactions at Interfaces. *J. Am. Chem. Soc.* **2004**, *126* (21), 6784–6797. https://doi.org/10.1021/ja049085k.
(13) Netz, R. R.; Andelman, D. Neutral and Charged Polymers at Interfaces. *Phys. Rep.* **2003**, *380* (1), 1–95. https://doi.org/10.1016/S0370-1573(03)00118-2.
(14) Murphy, M. C.; Rasnik, I.; Cheng, W.; Lohman, T. M.; Ha, T. Probing Single-Stranded DNA Conformational Flexibility Using Fluorescence Spectroscopy. *Biophys. J.* **2004**, *86* (4), 2530–2537. https://doi.org/10.1016/S0006-3495(04)74308-8.
